# Chemically Engineered Carbon Nanotubes Map Class-Selective Metabolite Enrichment from Human Plasma

**DOI:** 10.64898/2026.06.02.729405

**Authors:** In-Jun Hwang, Jada S. Gray, Mijin Kim

## Abstract

The spontaneous adsorption of biomolecules onto nanoparticle surfaces has been extensively characterized at the protein level, but the metabolite corona remains poorly defined while being physicochemically and biologically distinctive. Herein, we report the first class-level mapping of metabolite corona composition across 25 chemically modified carbon nanotubes in human plasma using untargeted liquid chromatography-mass spectrometry. Complementary analytical conditions detected approximately 9,000 metabolite features, of which over 5,000 yielded valid corona-versus-plasma enrichment measurements. Machine learning classifiers extended metabolite class annotations from 10-45% to the full detected set, enabling systematic analysis of class-level enrichment patterns. We find that polymer wrapping dominates corona composition, with DNA wrapping selectively enriching nonpolar lipids and PEG wrapping favoring polar metabolites. Within each polymer background, covalent quantum well defects further modulate class-level enrichment in a structure- and chemistry-dependent manner. Carboxyl aryl defects broadly enhance amphiphilic lipid recruitment, while trifluoro aryl defects suppress single-chain amphiphilic species but attenuate depletion of double-chain phospholipids. Our findings demonstrate that engineered nanotubes can serve as chemically tunable, selective scaffolds for metabolite enrichment, potentially enhancing the capability of nanotube-based platforms to recruit and detect structurally diverse, low-abundance small molecules in complex biofluids.

## Introduction

When nanomaterials enter a biological system, a layer of biomolecules spontaneously adsorbs to the surface, forming a biocorona. This interface is dynamic and competitive, shaped by differential adsorption affinities and exchange kinetics.^1, 2^ The biocorona confers a distinct biological identity on nanoparticles and has important implications for the physiological states of biological systems.^3^ The potential of the protein corona has been extensively investigated in the context of cellular uptake, trafficking,^4^ drug delivery,^5^ and biomarker discovery.^6–8^ However, the protein-centric view captures only part of the picture. Plasma contains thousands of lipid species, nucleotides, and small molecule metabolites, such as amino acids, organic acids, acylcarnitine, and benzenoids, all of which are available to adsorb competitively with proteins onto nanoparticle surfaces.^9^

Yet the metabolite corona has received far less attention than its protein counterpart, despite the fact that metabolites are the closest molecular readouts of disease phenotypes.^10^ The metabolome captures the integrated output of the genome, transcriptome, proteome, and environmental exposure in real time, and thus reflects the dynamics of biological activity, including enzyme function, energy flux, and the levels of signaling molecules that mediate intracellular communication.^11^ The selective adsorption of metabolites onto a nanoparticle surface could thus encode important clinical and biological information from the sample.^3^ Identifying the key chemical parameters that determine this selectivity would enable rational control over metabolite, lipid, and protein compositions in the biocorona.^3, 12^ However, these parameters remain poorly defined. The broad physicochemical diversity of metabolome requires multiple extraction solvents, chromatographic modes, and ionization polarities, introducing batch effects that confound quantitative comparison across nanoparticles. Low annotation rates inherent in untargeted LC-MS/MS data prevent a holistic view of enrichment patterns at the metabolite class level. Addressing these challenges requires a nanoparticle platform with independently variable surface chemistry, paired with an analytical framework that integrates multi-modal extraction data and extends class-level annotation to the dark metabolome.

Single-walled carbon nanotubes (CNTs) are well-suited as a model system for this investigation. Their one-dimensional cylindrical structure, composed of sp^2^-hybridized carbon atoms, can be tuned through polymer wrapping^13^ and quantum well defects (QWDs).^14^ Non-covalent wrapping with single-stranded DNA oligonucleotides or polyethylene glycol (PEG) defines the surface polarity and charge and creates specific binding interactions on the nanotube sidewall.^15–17^ Covalent functionalization of sp^3^ functional groups can create localized QWDs whose hydrophobicity, charge, and electronic character can be tuned by their terminating groups.^14^ This two-dimensional design space enables diverse variation of surface chemistry. In addition, semiconducting carbon nanotubes exhibit intrinsic near-infrared fluorescence^18^ that is sensitive to the adsorbed biomolecules.^6, 19^ The combination of a chemically programmable surface and an environmentally sensitive fluorescence response makes carbon nanotubes a unique platform to study how surface modifications shape metabolite corona composition.

Herein, we report the first systematic characterization of the metabolite corona on chemically modified carbon nanotubes. The nanotube library was constructed by five covalent surface modifications of nanotubes with QWDs and non-covalent polymer coatings with ssDNA and lipid-PEG. Metabolite corona analysis was conducted using untargeted LC-MS/MS across complementary analytical conditions to cover metabolite diversity from nonpolar lipids to polar small molecules. To extend class-level analysis to the majority of detected features, we developed machine-learning models that predict class labels of unannotated features using instrument-level descriptors. The optimized random forest classifiers accurately predicted nonpolar lipid classes but exhibited moderate and low prediction accuracy for structurally diverse polar classes. We employed a tiered analytical framework to handle the range of metabolite classes and found that polymer wrapping type sets the overall polarity profile. DNA wrapping selectively enriches nonpolar lipids while PEG wrapping favors polar metabolites. QWD chemistry tunes class-level enrichment in a polymer-dependent manner. Specifically, carboxyl aryl QWDs broadly enhance the adsorption of amphiphilic lipids, whereas trifluoro aryl QWDs exhibit structural selectivity that depletes single-chain amphiphilic species while attenuating the depletion of double-chain phospholipids. Our findings provide mechanistic insight into how surface chemistry shapes the identity of the metabolite corona, with direct implications for rational biocorona engineering and nanosensor design, and biomarker discovery.

## Results

### Construction and characterization of a polymer-wrapped QWNT library

To investigate how surface chemistry of carbon nanotubes shapes the small-molecule corona, we constructed a nanotube library comprising four covalent QWD variants and one pristine condition, each wrapped with five distinct polymers (**Fig 1a**). Covalent functionalization of sp^3^ defects created four QWNTs, including 4-carboxyl (carboxyl), 3,5-dinitro (dinitro), 3,4,5-trifluoro (trifluoro), and *N,N*-diethyl-4-amino (amino) aryl QWD functionalized nanotubes. The QWDs span a 5.7 log-unit hydrophobicity range (logP) and a charge-state range at pH 7.4 from −0.999 to +0.126 (Supplementary Table 1). (6,5)-enriched fractions were then isolated from mixed chirality QWNTs using aqueous two-phase extraction^20^ (see Methods). The enrichment of (6,5) chirality (>85%) was confirmed based on the characteristic E_11_ absorbance peak at 985 nm with minimal (6,4) nanotubes at 880 nm (**Fig 1b**; Supplementary Fig 1). Successful QWD formation was characterized by the emergence of a red-shifted E_11_^-^ emission peak in near-infrared fluorescence spectra (**Fig 1c**; Supplementary Fig 2) and by the increased Raman D band-to-G band ratios from 0.04 (pristine) to 0.15–0.18 across all QWNTs^21^ (**Fig 1d**; Supplementary Fig 3). Each QWNT and pristine nanotube was independently wrapped with three single-stranded DNA oligomers, (GT)_15_, (AT)_15_, and CT_2_C_3_T_2_C, or two PEG-lipid amphiphiles, DSPE-PEG(2000)-amine and DMG-PEG(2000), generating 20 QWNTs and 5 pristine nanotubes. Zeta potential measurements of the final QWNT suspensions confirmed that DNA-wrapped QWNTs retained a strong negative surface potential, while PEG-wrapped QWNTs were slightly positive or negative in buffer (**Fig 1e**).

**Figure 1.**
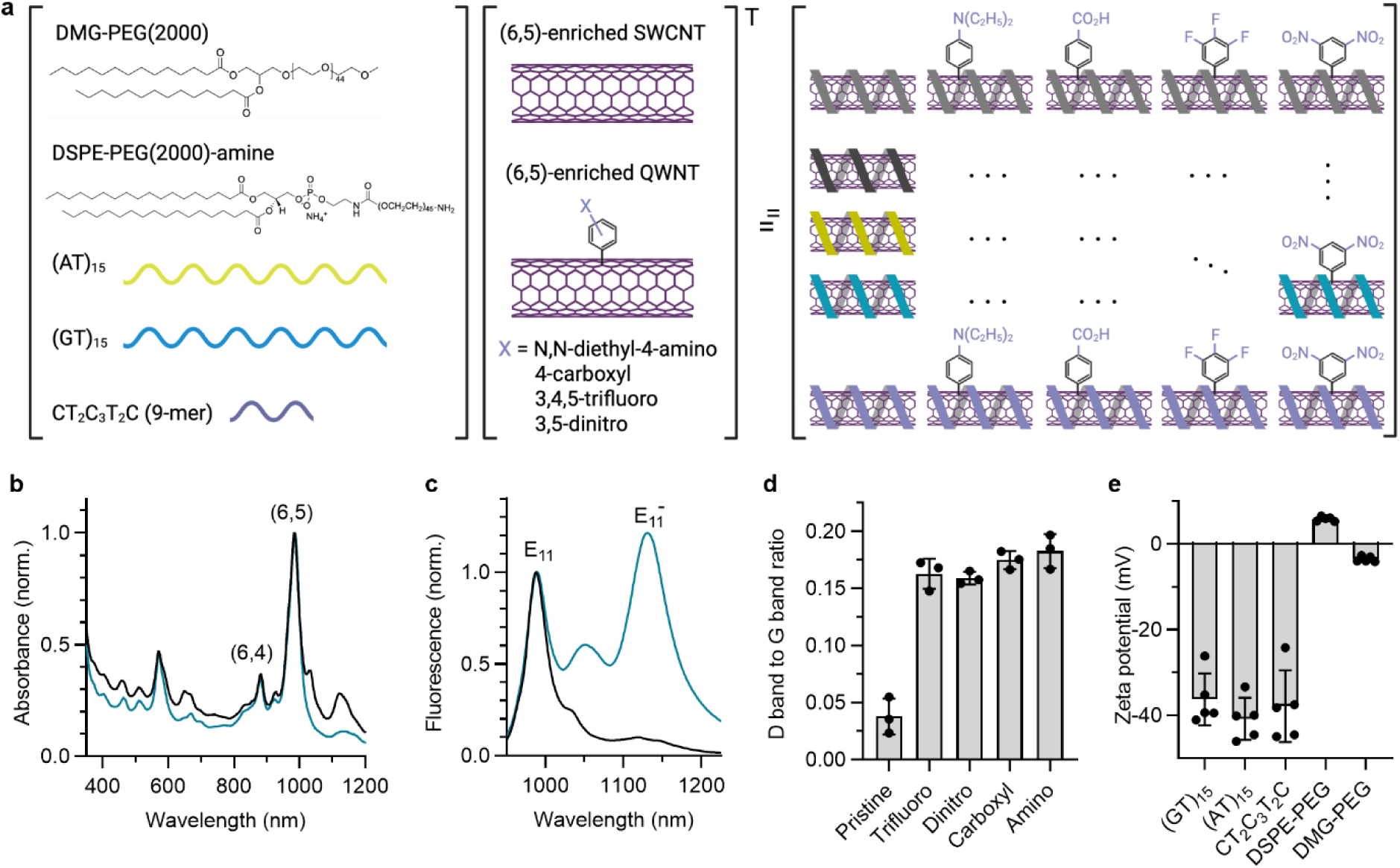
Polymer-QWNT library for metabolite corona analysis. **a,** Schematic of the library design. **b,** Absorption spectra of *N,N*-diethyl-4-aminoaryl functionalized carbon nanotubes in 1% sodium deoxycholate (aq) before and after aqueous two-phase extraction (black and blue, respectively). **c,** Normalized fluorescence spectra of purified pristine (black) and *N,N*-diethyl-4-aminoaryl functionalized (6,5)-enriched nanotubes (blue) in 1% sodium deoxycholate (aq), excited at 577 nm. **d,** Raman D to G ratio of (6,5)-enriched pristine nanotubes and QWNTs. **e,** Zeta potential of *N,N*-diethyl-4-aminoaryl functionalized (6,5)-enriched nanotubes with different polymer wrappings. Error bars represent the standard deviation of technical replicates.

### Multi-modal extraction workflow enables broad-coverage metabolite corona profiling

The QWNTs were incubated in 20% v/v of pooled human plasma at a nanotube concentration of 8 µg mL^-1^ (**Fig 2a**). After a 2-hour incubation, the mixture was centrifuged to pellet the corona, and the pellet was washed 3 times with phosphate-buffered saline to remove unbound biomolecules. To recover both the nonpolar lipid-rich fraction and the polar small-molecule fraction of the corona, we evaluated five extraction solvents spanning a polarity range from biphasic chloroform:methanol:water to monophasic acetonitrile:methanol (**Fig 2a**, see Methods). The eluents were analyzed using positive and negative ionization across reverse-phase (RP) and hydrophilic interaction (HILIC) LC-MS/MS columns, yielding four analytical modes per condition (**Fig 2b**, **Table 1**). Note that all samples underwent RP chromatography. Additionally, the extracts exclusively compatible with hydrophilic interaction were analyzed on a ZIC-HILIC column.^22^

**Figure 2.**
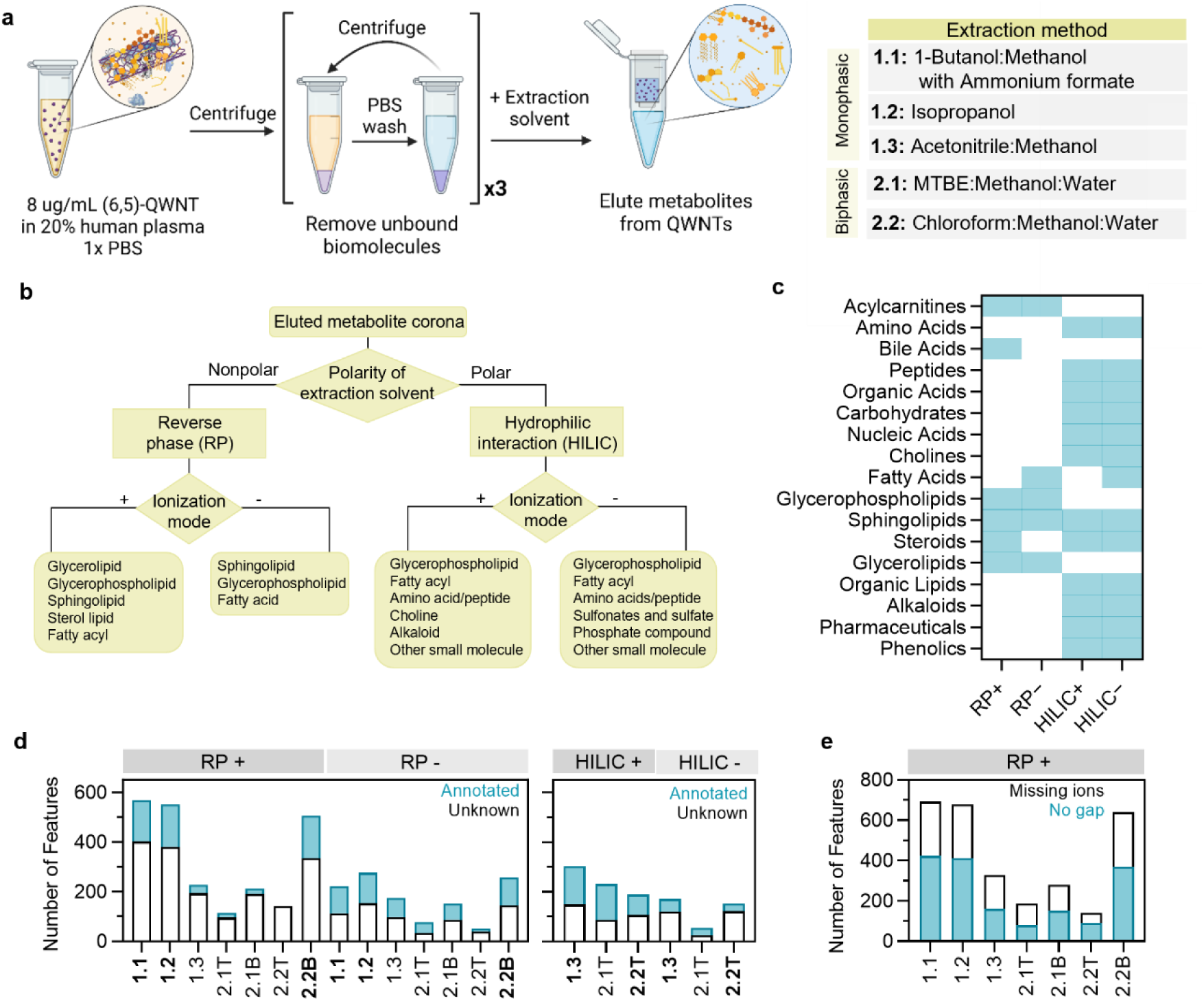
Metabolite extraction workflow and method optimization for corona profiling using (AT)_15_ wrapped, *N,N*-diethyl-4-aminoaryl functionalized (6,5) carbon nanotubes. **a,** Experimental workflow for metabolite extraction from carbon nanotubes in plasma. **b,** Diverse metabolite classes detected by varying the extraction solvent polarity, chromatography column, and ionization mode. **c,** Binary heatmap of metabolite class coverage across column and ionization mode combinations, where blue indicates classes detected in each LC-MS/MS mode. **d,** Feature counts of annotated (blue) and unknown (white) per extraction method and LC-MS/MS mode; extraction methods highlighted in bold are the down-selected methods for downstream analyses. **e,** Feature counts of no gap (blue) and missing ions (white) in selected extraction methods (RP+ mode).

**Table 1.** Formulation of extraction solvents for method validation. RP: Reverse phase (C30 column), HILIC: Hydrophilic interaction (ZIC^®^-HILIC column)

| Method | Formulation | Extraction Type | Chromatography compatibility | References |
| --- | --- | --- | --- | --- |
| 1.1 | 1:1 (v:v) 1-butanol: methanol with 5 mM ammonium formate | Single | RP | <sup>44</sup> |
| 1.2 | Isopropanol | Single | RP | <sup>45</sup> |
| 1.3 | 1:1 (v:v) methanol: acetonitrile | Single | HILIC | <sup>45</sup> |
| 2.1 | 2.6:2:2.4 (v:v:v) methyl-tert butyl ether: methanol: water | Biphasic | Top phase: HILIC<br>Bottom phase: RP | <sup>46</sup> |
| 2.2 | 2:2:1 (v:v:v) Chloroform: methanol: water | Biphasic | Top phase: HILIC<br>Bottom phase: RP | <sup>47</sup> |

To determine which combination of extraction solvent, chromatography mode, and ionization polarity most comprehensively captures the metabolite corona, we evaluated feature coverage and yield across all four analytical modes (RP+, RP-, HILIC+, HILIC-). As a model system, (AT)_15_-wrapped, *N,N*-diethyl-4-amino aryl QWNTs were used. Binary heatmap analysis confirmed that no single analytical mode captured the full breadth of corona-associated chemical classes (**Fig 2c**). Polar metabolite classes were more consistently detected in HILIC modes, whereas nonpolar lipid species were preferentially captured in RP modes, suggesting that orthogonal chromatographic coverage is essential to minimize profiling bias.

Extraction methods were then down-selected using a sequential two-criterion decision flow (Supplementary Fig 4). Methods with more than 100 annotated features and covering at least 6 (RP) and 9 (HILIC) distinct metabolite classes were retained for downstream analysis. Methods 1.1, 1.2, and 2.2B for RP, and 1.3 and 2.2T in HILIC modes passed the screening criteria (**Fig 2d**). Within these five down-selected methods, the number of no gap and missing ion features was over 139 across all selected conditions (**Fig 2e**, Supplementary Fig 5), indicating that the optimized extraction workflow reliably recovers a substantial metabolite feature set. Analytical reproducibility was confirmed across all 25 nanotube conditions, with a median coefficient of variation (CoV) of 20% in technical triplicates (Supplementary Fig. 6). Over 85% of nanotube method combinations exhibited CoV below 30%, consistent with standard reproducibility thresholds for untargeted metabolomics quality control.^23^ Batch-to-batch reproducibility was assessed using three independent nanotube batches based on CoV, relative log expression, and PCA of log_2_ intensities (see Methods and Supplementary Fig 7). The inter-batch CoV is within acceptable limits for untargeted metabolomics (23-37%).^23^ Consistent intensity distributions across batches in relative log expression analysis indicate the absence of systematic batch variation. Modest inter-batch separation in PCA reflects differences in the subset of features detected above the signal threshold in each batch. These results confirm that downstream enrichment analyses were not confounded by technical variation between experimental runs. Based on the optimized extraction and LC-MS/MS workflow, we extend the metabolite corona analyses to the remaining 24 nanotube formulations.

To integrate metabolite abundance data across the five extraction methods (**Table 1**) while accounting for method-level batch effects, we applied an inverse-variance-weighted (IVW) meta-analysis^24, 25^ (see Methods). Briefly, each extraction method was treated as an independent measurement, and the method-specific log_2_ fold changes (log_2_FC) were combined using precision weights (1/SE^2^) so that more consistent measurements contributed proportionally more to the final estimate (Eq 1).

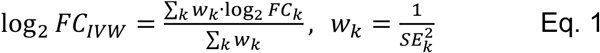

wherein w_k_ and SE_k_ are the precision weight and standard errors of the extraction method *k*. The IVW-combined log_2_FC and p-values were computed for each feature in two comparisons: 1) All nanotube coronas relative to plasma and 2) each QWNT corona relative to its pristine baseline within each polymer wrapping (Supplementary Table 2).

### Machine learning classifiers expand metabolite class annotation to the full feature set

Spectral library annotation implemented in Compound Discoverer identified 677 RP features and 1,081 HILIC features with varying class assignments, leaving 6,486 RP features and 1,686 HILIC features unannotated (**Fig 2d**). To enable class-level enrichment analysis across the entire detected feature set, we trained supervised machine learning models using three instrument-level descriptors for each feature: mass-to-charge ratio (m/z) for molecular weight, mass defect for elemental composition, and ion polarity for charge-carrying functional groups (**Fig 3a**). Three classifiers (k-nearest neighbors, KNN; random forest, RF; gradient boosting, XGBoost) were trained with 3-fold cross-validation on 80% of annotated data and evaluated on a held-out 20% test set, stratified by class (see Methods). To improve model performance in multiclass classification, we consolidated the metabolite classes into five and nine classes for RP and HILIC, respectively (Supplementary Table 3). The consolidation decisions are grounded in the physical chemistry of MS, chromatographic retention behavior, and the known biosynthetic relationships between metabolite classes.

**Figure 3.**
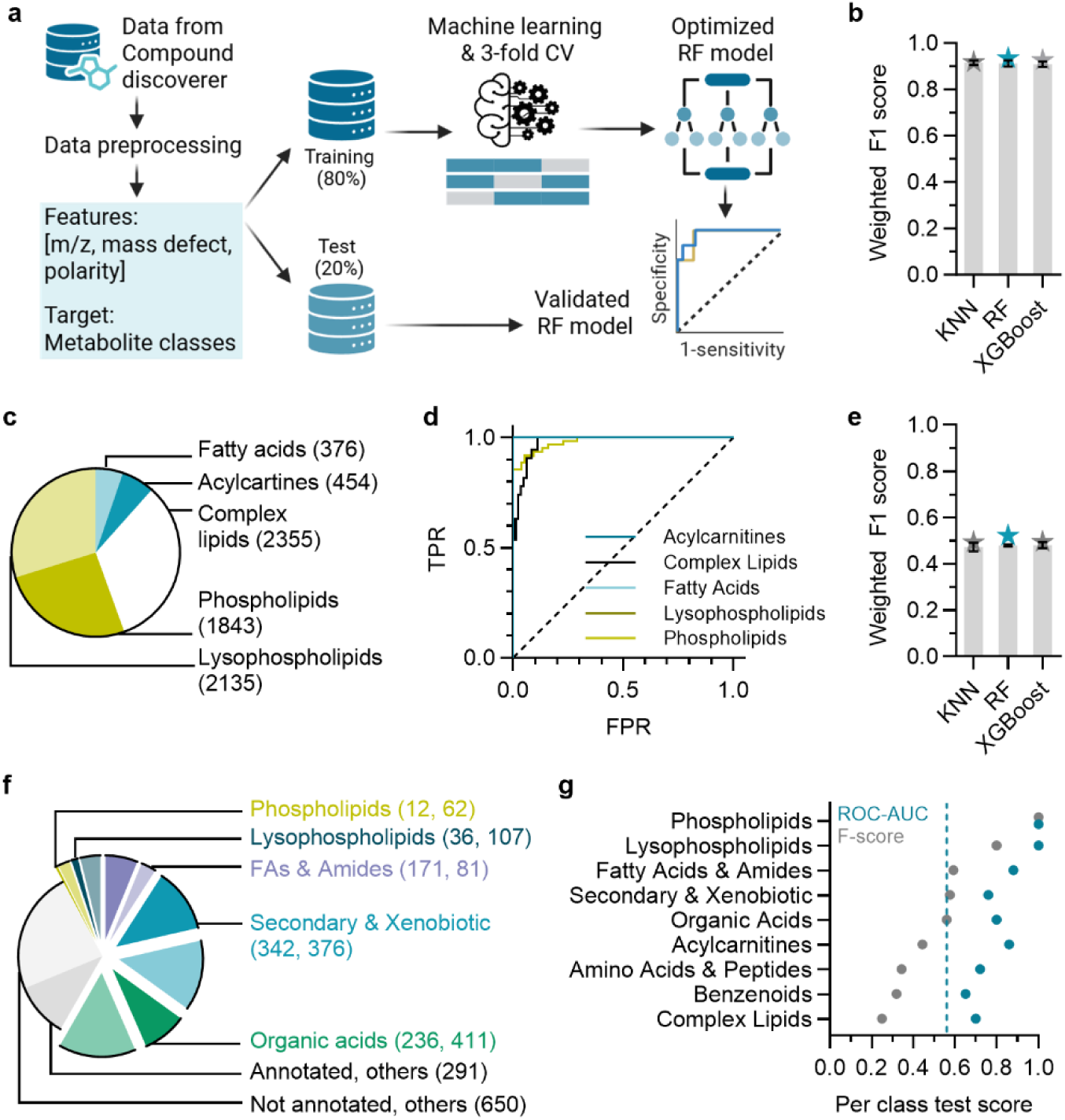
Machine learning classification of unannotated corona features into metabolite classes. **a,** Classification pipeline. Features exported from Compound Discoverer were characterized by m/z, mass defect, and ESI polarity. Annotated features (677 RP; 1,081 HILIC) are used for prediction model development. Three classifiers (k-nearest neighbors, KNN; random forest, RF; XGBoost) were trained with 3-fold cross-validation on 80% of annotated data and evaluated on a held-out 20% test set. **b,** Weighted F1 scores (test set) for RP mode. **c,** Class distribution of RP-annotated and predicted features. **d,** Per-class test ROC curves of the best-performing multi-class classification model for RP mode. **e,** Weighted F1 scores (test set) for HILIC mode. **f,** Class distribution of HILIC-annotated (solid) and predicted (semi-transparent) features. The remaining four minority classes are aggregated in the Others category. **g,** Per-class test set score for HILIC mode classification. The vertical dashed line indicates the F1-score threshold for high-confidence classes. For **b** and **e**, error bars show standard deviation across cross-validation folds and star symbols indicate test set scores.

For the RP feature set, the optimized multi-class RF model achieved a weighted F1 score of 0.934, which falls within the standard deviation of the 3-fold cross-validation scores (**Fig 3b**, Supplementary Fig. 8, Supplementary Table 4). The high prediction accuracy reflects the well-separated distributions of lipid classes in the m/z and mass-defect feature spaces (Supplementary Fig. 9). Complex lipids and phospholipids occupy distinct high m/z ranges, acylcarnitines feature low mass defects, and lysophospholipids are separable from phospholipids by mass defect at overlapping m/z values. When applied to the full RP feature set, the RF classifier assigned metabolite class labels to all detected features (**Fig 3c**). The ROC-AUC for all five classes, including phospholipids, complex lipids, lysophospholipids, acylcarnitines, and fatty acids, achieved over 0.98 (**Fig 3d**, Supplementary Fig. 10, Supplementary Table 4).

For the HILIC feature set, nine-class classifiers were trained on a structurally more diverse set of polar and lipid-adjacent classes. The best-performing model was RF, with a weighted F1 score of 0.525 on the held-out test set (**Fig 3e**, Supplementary Fig. 11, Supplementary Table 4). The model performance was substantially lower than that of the RP classifiers, and 19.7 % of features did not meet the confidence threshold and were classified as low confidence features (**Fig 3f**). The modest performance in HILIC is attributed to greater spectral overlap among polar metabolite classes, a larger number of metabolite classes, and a smaller training set per class (Supplementary Fig 12). Per-class test scores varied across the nine HILIC classes, ranging from 0.750 for lysophospholipids to 0.286 for complex lipids, with per-class ROC-AUC ranging from 0.66 for benzenoids to 0.99 for lysophospholipids (**Fig 3g**, Supplementary Fig 13). Because reliable class-level enrichment analysis requires that class assignment be sufficiently accurate, we implemented a tiered analytical strategy based on individual per-class F1 performance and the number of annotated features per class.

Classes with a per-class F1 score of ≥ 0.56. Track 1 includes all metabolite classes of RP and phospholipids, lysophospholipids, organic acids, fatty acids & amides, and secondary & xenobiotic compounds for HILIC. For the remaining HILIC classes, we analyzed the metabolite enrichment patterns for annotated and predicted features separately and compared their consistency for downstream analyses (Track 2). This tiered framework ensures that all HILIC features are included in enrichment analyses while maintaining statistical validity within each tier.

### Nanotube corona selectively enriches differential metabolite classes relative to plasma

Prior to class-level enrichment analysis, we applied a deduplication step to collapse redundant adduct and in-source fragment features of the same metabolite into a single representative entry (see Methods), reducing the feature count to 6,383 (RP) and 2,529 (HILIC). Acylcarnitines and amino acids & peptides in HILIC were most affected due to the prevalence of sodium and potassium adducts in these classes (Supplementary Table 5). All class-level enrichment analyses, volcano plots, and pathway analyses reported below use the deduplicated feature set. ML classification and IVW fold-change estimation were performed on the full feature set prior to deduplication. A sensitivity analysis on the full feature set confirmed that all class-level conclusions were preserved (Supplementary Table 6).

To establish the baseline corona composition and distinguish selective adsorption, metabolite profiles from corona eluates were compared with those from matched human plasma. The corona detected a higher number of no-gap features than plasma in RP, whereas HILIC modes detected the opposite trend (**Fig 4a**). Missing ion features were consistently lower in the corona than in the plasma across all four modes. We restricted the downstream analysis to no gap and missing ions features, for which standard gap-filling imputation by Compound Discoverer reliably recovers intensity values from raw spectral data. Feature overlap analysis revealed that a substantial proportion of corona features were uniquely detected in the corona, with HILIC modes showing a larger corona-unique fraction compared to RP modes (**Fig 4b**).

**Figure 4.**
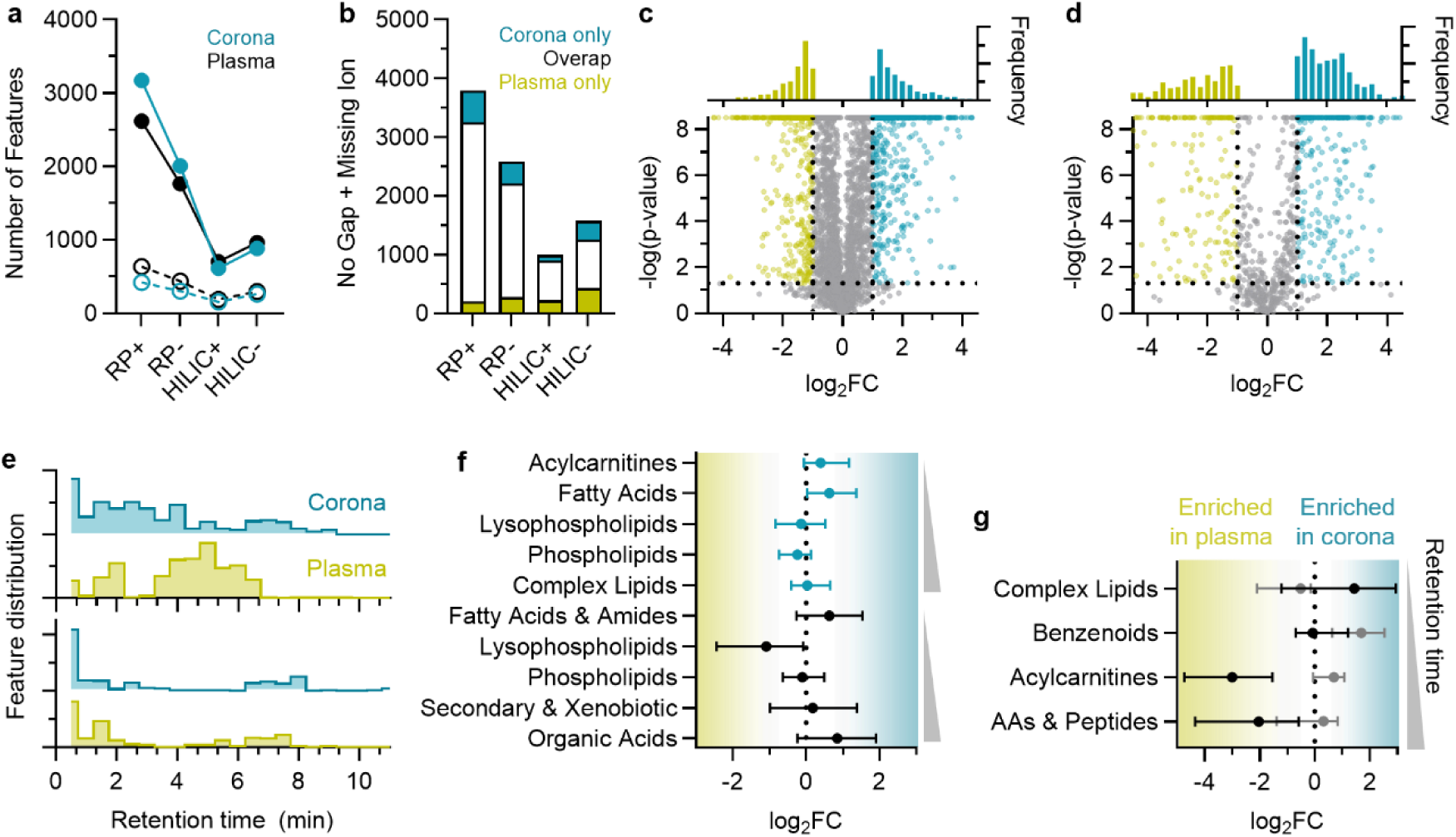
Global metabolite corona composition relative to plasma. **a,** The number of no gap features (filled circles) and missing ion features (open circles) in the full plasma extract (black) vs. corona extracts (blue) across all four LC-MS/MS modes. **b,** Stacked bar plot visualizing the number of no gap and missing ions features unique to the metabolite corona (blue) and plasma (yellow), as well as the number of overlapped features (white) between corona and plasma across all modes. Volcano plots comparing nanotube corona to plasma for **c,** RP and **d,** HILIC. Log_2_FC represents corona enrichment minus plasma enrichment. Blue and yellow points represent corona-enriched and plasma-enriched features, respectively, above the significance thresholds (p-value cutoff of 0.05 and |log_2_FC| cutoff of 1, dashed lines). The top panels represent the frequency distribution of significant features. **e,** Retention time distribution analysis of significant features detected in the metabolite corona and plasma in RP (top blue) and HILIC (bottom black). **f,** Class-level enrichment for corona vs. comparisons, showing RP classes (top) and HILIC Track 1 classes (bottom), ordered by median retention time. Positive and negative log_2_FC indicate metabolite classes enriched in the corona and in plasma, respectively. **g,** Class-level enrichment for HILIC Track 2, ordered by median retention time. Circle symbols indicate the median of log_2_FC of annotated (black) and predicted features (gray). Horizontal bars indicate the interquartile range (25-75%) across annotated and predicted features.

Volcano plot analysis of IVW-combined log_2_FC revealed that corona-enriched and plasma-enriched features were distributed asymmetrically across RP and HILIC modes. Of 3,943 RP features analyzed, 1,098 (28%) were significantly enriched or depleted in the corona versus plasma (p<0.05, |log_2_FC|>1 (**Fig 4c**). Of 1,037 HILIC features, 478 (46%) showed significant enrichment or depletion (**Fig 4d**), indicating that the nanotube corona is a highly selective subset of the plasma metabolome. Retention time distribution analysis of the significant features (p<0.05 and log_2_FC >1) showed that corona-enriched features in RP were distributed across both early and later retention times, while plasma-enriched features were concentrated in the middle retention time range (**Fig 4e**, top, Supplementary Fig 14). In HILIC, both corona and plasma-enriched features were concentrated at earlier retention times (**Fig 4e**, bottom, Supplementary Fig. 15). These differential elution patterns indicate that the nanotube surface simultaneously recruits nonpolar lipids and polar small molecules (Supplementary Fig 16).

Class-level enrichment analysis using median IVW-combined log_2_FC confirmed that the corona selectively recruits specific metabolite classes relative to plasma (**Fig 4f**). In RP, acylcarnitines and fatty acids were enriched in the corona, consistent with their early elution in the RP column in **Fig 4e**, while phospholipids and lysophospholipids were more abundant in plasma. Complex lipids were near no change. In HILIC Track 1, organic acids, fatty acids & amides, and secondary & xenobiotic metabolites were modestly enriched in the corona, whereas phospholipids and lysophospholipids were more abundant in plasma. Consistently, the proportion of features individually significant in the enriched versus depleted directions was clearly dominated by the class-level trend (Supplementary Fig 17).

For HILIC Track 2 classes, the full distribution of per-feature log_2_FC for each class, split by annotated vs. predicted, visualizes the ensemble-level enrichment patterns with respect to plasma (**Fig 4g**). All Track 2 classes showed the opposite enrichment trend between the annotated and predicted subsets, likely reflecting misclassification of polar corona-enriched features. For these classes, we defer interpretation to the polymer- and QWD-level analyses in the following sections, which assess enrichment rank distributions without relying on individual class-label accuracy.

### Polymer wrapping significantly modulates the corona composition

Pristine nanotubes showed selective metabolite enrichment relative to plasma across all five polymer wrappings (Supplementary Fig. 18). Principal component analysis (PCA) of RP corona profiles revealed that polymer identity explained 43.2% of total variance on PC1 for pristine nanotubes, with PEG-wrapped nanotubes (DMG, DSPE) separating cleanly from DNA-wrapped nanotubes ((GT)_15_, (AT)_15_, CT_2_C_3_T_2_C) along this axis (**Fig 5a**, Supplementary Fig 19). When QWNTs were included, polymer class remained the dominant source of variance while defect identity produced scatter within each polymer cluster (**Fig 5b**; Supplementary Fig 20). To assess polymer effects exclusively, we focused our analysis on pristine nanotubes (**Figs 5c-f**).

**Figure 5.**
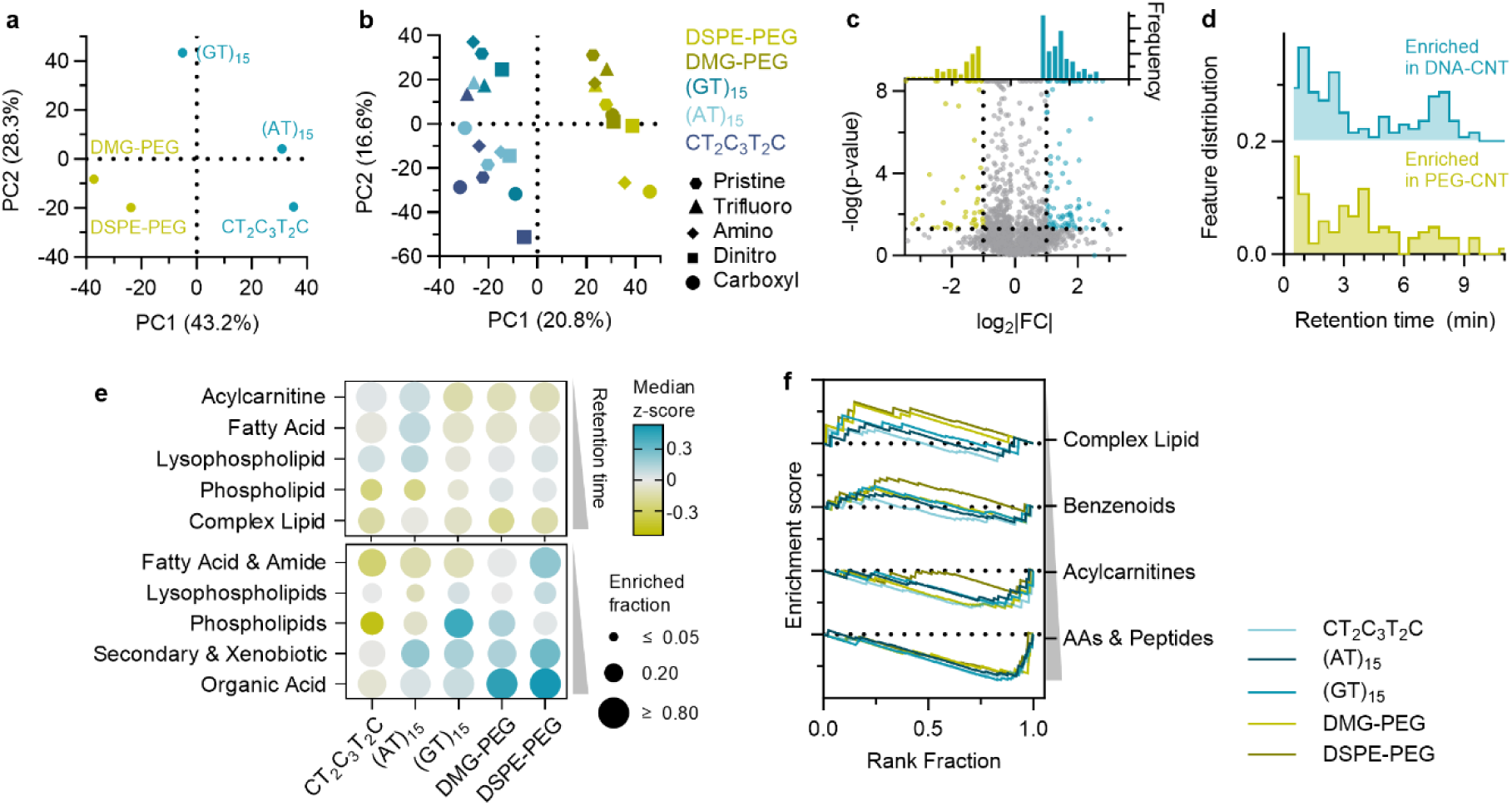
Effect of polymer wrapping type on metabolite corona composition. **a,** Principal component analysis (PCA) of corona profiles across the five polymer wrappings (pristine nanotubes only) in RP. **b,** PCA of corona profiles across all 25 nanotube combinations (20 polymer-QWNTs + 5 pristine nanotubes) in RP. Marker shape and color indicate QWD type and polymer type, respectively. **c,** Volcano plot comparing coronas extracted from DNA-wrapped to PEG-wrapped pristine nanotubes (RP). Log_2_FC represents IVW-DNA corona enrichment minus IVW-PEG corona enrichment. P-value from Welch’s t-test. Blue and yellow points are features significantly enriched in DNA-CNT and PEG-CNT coronas, respectively (p <0.05, |log_2_FC|>1). The top panel shows the frequency distributions of significant features. **d,** Retention time distributions of significant features detected in **c**. **e,** Class-level enrichment bubble plot for the DNA vs PEG comparison (pristine only), showing RP classes (top) and HILIC Track 1 classes (bottom). Dot size indicates the enriched fraction, and color indicates the median z-score of log_2_FC normalized across polymers within each class. **f,** Running enrichment scores for HILIC Track 2 metabolite classes across polymer wrapping types. Features are ranked by log_2_FC (corona vs. plasma) within each polymer wrapping. Monotonically decreasing curves indicate that class members are depleted from the nanotube corona relative to plasma; increasing curves indicate corona enrichment. Dotted lines denote the zero-enrichment baseline.

In RP mode, pairwise comparison of DNA- versus PEG-wrapping identified that 149 features (6.2% of the total identified features) were significantly different (**Fig 5c**). When significant features with |log_2_FC| of higher than 1 were analyzed, features enriched in DNA-CNT coronas clustered at early (1.0-2.5 min) and late (7.0-8.0 min) retention time (**Fig 5d**, top). Features enriched in PEG-wrapped nanotube coronas clustered at shorter (0.5-1.0 min) and mid (3.5-4.5 min) retention times (**Fig 5d**, bottom). Similarly, we observed polymer class dependent enrichment and retention time in HILIC mode (Supplementary Fig. 21).

Class-level enrichment analysis across different polymer wrapping types provides mechanistic insights into CNT-metabolite interactions (**Fig 5e**, Supplementary Fig 22). In RP, fatty acids (titratable anionic, amphiphilic) and lysophospholipids (single chain, moderately hydrophobic) were enriched on (AT)_15_ and CT_2_C_3_T_2_C wrapped nanotubes. The preferential enrichment is presumably due to the fact that DNA wrapping exposes the hydrophobic sidewall of nanotubes,^26–29^ enabling hydrophobic contact with lipid acyl chains. Acylcarnitines with a permanent positive charge at pH 7.4 also showed sequence-dependent enrichment among DNA-CNT coronas. (AT)_15_wrapping showed the strongest enrichment, driven by electrostatic interactions with the negatively charged DNA backbone, modulated by DNA stacking geometry and stability.^29^ Phospholipids were plasma-enriched across all coronas but less depleted in PEG-CNTs, while complex lipids showed no distinct preference in the class median. In HILIC Track 1, organic acids were enriched in PEG-CNT coronas, presumably due to the positive interactions between hydrophilic PEG wrapping and polar metabolites.

To assess Track 2 classes without relying on individual class-label accuracy, we ranked all HILIC features by their log₂FC (corona vs plasma) within each polymer wrapping and examined where members of each Track 2 class fell within this ranked list (**Fig 5f**, Supplementary Table 7). If class members are concentrated among the most corona-enriched features, the running enrichment score (ES) rises. Conversely, if concentrated among the most plasma-enriched, ES falls. Complex lipids showed positive enrichment scores across all five polymers, indicating consistent corona enrichment, with PEG-wrapped nanotubes showing the strongest enrichment. Acylcarnitines, and amino acids & peptides showed uniformly negative enrichment scores across all polymers, indicating depletion from the corona relative to plasma. Benzenoids showed a mixed pattern, with modest enrichment in PEG-wrapped CNTs and depletion in DNA-wrapped CNTs.

### QWD modulates corona composition through hydrophobicity matching

To isolate QWD chemistry contributions from the polymer-wrapping background, all QWNT conditions were compared against their matched pristine-wrapped references. We quantified the fraction of features within each class that were significantly enriched (p < 0.05, |Δlog_2_FC| > 0.5) by at least one of the 20 polymer-QWNT combinations (**Fig 6a**). Across all RP features, fatty acids (40%) and acylcarnitines (38%) showed the highest sensitivity to QWD introduction (either depleted or enriched), followed by complex lipids (35%) and lysophospholipids (29%), with phospholipids the lowest (22%). Note that a substantial fraction of significant features were enriched in some polymer-QWNTs and depleted in others, suggesting the polymer dependent QWD effects discussed below.

**Figure 6.**
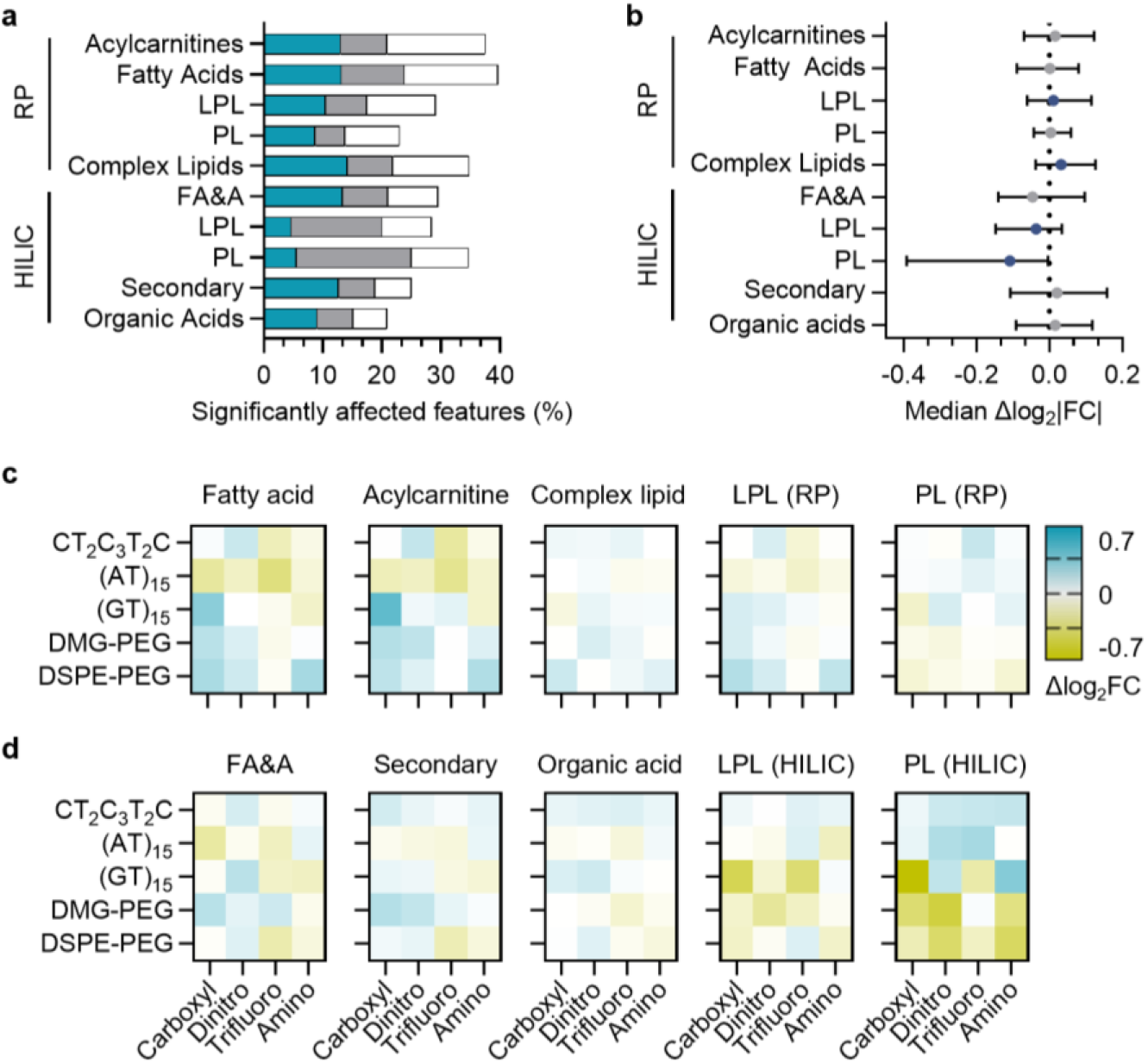
QWDs modulate class-selective metabolite enrichment in the QWNT corona. **a,** The percentage of features within each metabolite class that are significantly affected (p < 0.05, |Δlog_2_FC| > 0.5) by at least one of 20 polymer-QWNTs compared to the pristine control. Blue bars indicate the fraction of features exclusively enriched by QWDs; Gray bars indicate the fraction exclusively depleted; White bars indicate the fraction that is enriched in some polymer-QWNTs and depleted in others. Classes are shown for RP (top) and HILIC Track 1 (bottom), ordered by median retention time within each mode. **b,** Class-level differences in median Δlog_2_FC (all 20 QWNTs vs. pristine). Purple and gray circles indicate significant (Wilcoxon signed-rank test, Benjamini–Hochberg q<0.05) and not significant classes, respectively. Horizontal bars indicate the interquartile range (25-75%) across features. Polymer-QWD interaction heatmaps for Track 1 **c,** RP and **d,** HILIC metabolite classes across all nanotubes. Columns represent defect types ordered by polarity. Rows represent polymer wrappings. Color bar indicates the changes in the median log_2_FC (corona versus plasma) compared with pristine control. Positive values (blue) indicate enrichment by QWD; negative values (yellow) indicate depletion by QWD. LPL: Lysophospholipids. PL: Phospholipids. FA&A: Fatty Acids & Amides. Secondary: Secondary & Xenobiotic.

Class-level median enrichment scores revealed significant QWD effects for selected metabolite classes (**Fig 6b**). In RP, complex lipids (median Δlog_2_FC = +0.032) and lysophospholipids (+0.011) were significantly shifted by QWD introduction, whereas phospholipids, acylcarnitines, and fatty acids showed no significant class-level effect. In HILIC Track 1, phospholipids (–0.109) and lysophospholipids (–0.037) were significantly depleted by QWD introduction, while no other class reached significance. The opposing QWD response of RP and HILIC phospholipids and lysophospholipids reflects the distinct subpopulations captured by each chromatographic mode (Supplementary Fig 23).

Polymer-QWD interaction heatmaps of the median log_2_FC (QWNT vs pristine control) for all Track 1 metabolite classes across the 20 nanotube conditions revealed that pooled class-level statistics mask substantial polymer dependence (**Figs 6c-d**, Supplementary Fig 24). For most metabolite classes, individual polymer-QWD combinations produced effect sizes an order of magnitude larger than the pooled medians, with opposing directions across polymer wrappings. For instance, phospholipids detected in RP were significantly depleted in DSPE-PEG wrapped QWNTs but enriched in DNA-QWNTs (**Fig 6c**), which cancels at the aggregate level in **Fig 6b**.

Across all classes, the heatmaps show substantial polymer-QWD interactions. In RP, despite underlying pronounced polymer effects, two QWDs produced directionally consistent effects across most polymer backgrounds (**Fig 6c**). Carboxyl aryl QWDs produced net positive enrichment of amphiphilic lipid classes, including acylcarnitines, fatty acids, lysophospholipids, and complex lipids. The largest enhancements were observed on polymers with weaker pristine baselines, such as (GT)_15_, DMG-PEG, and DSPE-PEG, whereas the QWDs had minimal effect on (AT)_15,_ where pristine enrichment was already high. Trifluoro aryl QWDs depleted single-chain amphiphilic classes (acylcarnitines, fatty acids, and lysophospholipids) on polymers with a strong pristine baseline, (AT)_15_ and CT_2_C_3_T_2_C, while minimally changing the enrichment on other polymer-QWNTs. On the other hand, trifluoroaryl QWDs attenuated the depletion of double-chain phospholipids. Dinitro and amino QWDs produced weaker and less directional effects, with responses dominated by specific polymer-defect combinations.

In HILIC Track 1, QWD effects were generally larger in magnitude than in RP and exhibited mixed patterns across metabolite classes (**Fig. 6d**). Carboxyl and dinitro QWDs broadly enriched Secondary & Xenobiotic compounds across most polymer backgrounds. Dinitro QWDs uniquely enriched Fatty Acids & Amides, whereas trifluoro QWDs depleted this class. Lysophospholipids were depleted by carboxyl and dinitro QWDs across most polymer backgrounds. It showed reversed the enrichment pattern observed in RP. Phospholipids showed the largest polymer-dependent variation than that in RP. It is consistent with the wide interquartile range observed for phospholipids in **Fig. 6b**. The mechanism underlying these enrichment/depletion patterns remains unclear, but suggests that QWDs locally reorganize the polymer wrapping in QWD-specific ways, and modulate the hydrophilic interactions with polar metabolites.

Running enrichment score curves of HILIC Track 2 classes across the four QWDs were predominantly flat or irregular in trajectory (Supplementary Fig 25, Supplementary Table 8). This indicates that Track 2 class members are broadly distributed within the ranked feature list, reflecting the chemical heterogeneity within each broad class. However, acylcarnitines and benzenoids showed QWD-specific responses. Benzenoids were enriched by carboxyl and dinitro QWDs, while they were depleted with trifluoro and amino QWDs. Acylcarnitines showed a more complex defect response. Trifluoro, carboxyl, and dinitro QWDs all enhanced enrichment, while the amino defect uniquely reduced it.

Finally, we characterized the adsorbed metabolome at the level of biochemical pathways via mummichog pathway enrichment analysis^30^ of the HILIC dataset (Supplementary Fig 26, Supplementary Table 9). Lipid metabolism pathways, including de novo fatty acid biosynthesis, fatty acid activation, and the carnitine shuttle, were significantly over-represented among corona-enriched features. Interestingly, multiple amino acid metabolic pathways, including the urea cycle and tryptophan metabolism, etc., were also enriched despite the overall class-level depletion.

## Discussion

We developed a multimodal extraction workflow that captures the broad chemical diversity of the metabolite corona of chemically modified (6,5) enriched carbon nanotubes. By evaluating five extraction solvents across four LC-MS/MS analytical modes and applying a sequential selection rule, we identified a method combination that reliably recovers a reproducible, high-confidence set of metabolites. The IVW meta-analysis further suppresses method-level batch effects without distorting within-condition signals, establishing an analytical platform that could be extended to other nanomaterial systems. This multi-modal approach detected approximately 9,000 metabolite features and quantified corona enrichment for over 5,000 of these across the nanotube library, which are an order of magnitude larger in both feature count and condition breadth than previous nano-omics^31–34^ and five times larger than what is typically annotated in large-scale untargeted plasma metabolomics studies.^35, 36^

Machine learning the metabolite classes based on instrument-level descriptors (m/z, mass defect, ion polarity) enabled the classification of unannotated features. While the random forest classifier achieved near-perfect performance (weighted F1 score of 0.934, ROC-AUC of 0.98) for RP, the structurally diverse HILIC feature set required a tiered analytical strategy, in which high-confidence classes were analyzed by Wilcoxon rank-sum testing (Track 1), like RP features, and lower-confidence classes by enrichment score (Track 2). This tiered approach ensures that all detected features contribute to enrichment analyses while maintaining statistical validity appropriate to each class.

Comparison of the full nanotube corona against plasma confirmed that the corona is compositionally distinct, with selective enrichment of specific metabolite classes. Polymer- and QWD-level enrichment analyses indicate that a two-level hierarchy governs the metabolite corona on polymer-QWNTs. Polymer wrapping sets the overall corona polarity profile. DNA-wrapped nanotubes preferentially enriched amphiphilic single-chain lipid species, including fatty acids and lysophospholipids, through hydrophobic sidewall exposure and exhibited sequence dependent enrichment for cationic acylcarnitines. In contrast, PEG-wrapped nanotubes favored polar small molecules through their hydrophilic, sterically shielded surface. Within the polymer-defined baseline, covalent QWD chemistry further modulates enrichment of specific metabolite classes.

QWD effects are convoluted with the polymer effects, making it difficult to isolate the independent QWD contribution. The same QWD chemistry produces opposing enrichment patterns depending on the polymer wrapping, and no single QWD property, e.g, hydrophobicity, charge, or polarity, predicts the response across all polymer backgrounds. This coupling indicates that the defect site does not present an isolated chemical environment but rather a perturbation embedded within a polymer-defined surface architecture. Despite this complexity, two QWDs produced directionally consistent net effects on amphiphilic lipid classes. Carboxyl QWDs enhanced corona enrichment most strongly on polymers with weaker pristine baselines, while trifluoro QWDs suppressed enrichment on polymers with stronger baselines and modestly attenuated the depletion of double-chain phospholipids. This structural sensitivity suggests that QWD chemistry differentially affects binding interactions of single-/multi chain lipids on the nanotube sidewall.

Track 2 enrichment score analysis distinguished the adsorption patterns of benzenoids and acylcarnitines. Benzenoids showed enrichment with carboxyl and dinitro QWDs and depletion with trifluoro and amino QWDs. The enrichment pattern of benzenoids is consistent with preferential adsorption on polar QWDs. Positively charged acylcarnitines were uniquely attenuated by aminoaryl QWDs, presumably due to the local disruption of the negative surface charges by the positively charged QWDs. The sequence-dependent enrichment of acylcarnitines among negatively charged, DNA-wrapped nanotubes further suggests an electrostatic component to their adsorption.

Our resulting dataset, over 5,000 reproducibly quantified enrichment profiles mapped across 25 chemically engineered carbon nanotubes, constitutes the most comprehensive metabolite corona resource generated to date. The class-selective enrichment patterns reported here open several new directions for nanosensor design, biomarker discovery, and corona engineering in nanoparticles. The ability to preferentially enrich specific lipid classes through polymer-QWD selection provides a foundation for developing corona-phase molecular recognition-based nanosensors that target defined metabolite windows, e.g., monitoring specific lipid metabolites.^37, 38^ In addition, these engineered coronas could serve as nano-omics enrichment tools for biomarker discovery, selectively concentrating low-abundance metabolites or lipids from complex biofluids.^2, 3, 7^ If class-selective enrichment is preserved in plasma (e.g., from patients with lymphatic diseases, diabetes, or cancer), it may be possible to amplify disease-specific metabolite signatures that are diluted below detection limits in unfractionated plasma, thereby enabling a lipid-biopsy approach to early diagnosis.^6, 19^

On the other hand, several limitations of the current work suggest interesting questions for future investigation. First, while the present analysis resolves metabolite adsorption at the class level, intra-class structural heterogeneity, such as head-group identity, acyl chain length, and unsaturation, may further modulate nanotube chemistry defined affinity. Moreover, the non-monotonic grouping of QWD effects does not correlate with a single physicochemical descriptor, suggesting combinatorial effects of steric, electrostatic, and dispersion forces in metabolite-polymer-QWD interactions. Molecular dynamics simulations of small-molecule adsorption on carbon nanotubes would provide mechanistic insight into these subclass-level molecular interactions.^39, 40^ Second, while we demonstrate structure-driven correlations with biochemical signatures, mummichog analysis of an extended metabolomics dataset could reveal pathway-level selectivity to guide the rational design of machine perception lipid biopsies. Finally, investigating the formation dynamics and interactions between metabolites and proteins in biomolecular corona is critical for establishing a comprehensive multi-omics profile of the biomolecular interface on engineered nanomaterials.^12, 41^

## Methods

### Solid state synthesis of quantum well defect modified carbon nanotubes

QWNTs were synthesized using either peroxide-mediated aryl diazonium reduction or in situ diazonium formation.^42^ Briefly, amino and dinitro QWDs were prepared by reacting 2.0 mg of raw nanotubes (SG65i, Sigma Aldrich) with 4-(diethylamino)benzenediazonium tetrafluoroborate (2.0 mg mL^-1^) and 3,5-dinitrobenzenediazonium tetrafluoroborate (2.64 mg mL^-1^), respectively, followed by the addition of 200 and 197 μL of hydrogen peroxide (50% w/v, Fisher Scientific) and incubation in the dark for 48 h and 24 h, respectively. Trifluoro and carboxyl QWDs were synthesized via in situ diazonium formation using 3,4,5-trifluoroaniline (64 mM) and 4-aminobenzoic acid (84 mM) with NOBF_4_, respectively, followed by the addition of 200 μL of hydrogen peroxide (50% w/v, Fisher Scientific) and incubation in the dark for 24 h and 72 h, respectively, with 2.0 mg of raw nanotubes (SG65i, Sigma Aldrich) used for trifluoro and 1.0 mg for carboxyl. The QWNTs were thoroughly washed with water and ethanol by vacuum filtration on a 20 µm polyethylene frit.

### Chirality enrichment of QWNTs by aqueous two-phase extraction

1 mg of carbon nanotubes (either raw, unfunctionalized or QWD modified SG65i) was dispersed in 1 mL of aqueous solution of 1% (w/v) sodium deoxycholate (Sigma Aldrich). The mixture was sonicated using a probe-tip sonicator with 70% amplitude (FB120, Fisher Scientific) at 4 °C for 30 min. The resulting solution was centrifuged at 21,300 g for 30 min. The top 80% supernatant was collected. (6,5) Chirality enrichment was performed by aqueous two-phase extraction^20^. Briefly, 10.2 mL of Stock 1 solution containing sodium cholate (Sigma Aldrich), polyethylene glycol (6 kDa, Sigma Aldrich), sodium dodecyl sulfate (Sigma Aldrich), sodium chloride (Sigma Aldrich), and dextran (70 kDa, TCI Chemicals) was mixed with 699 µL of carbon nanotube suspension and 902 µL of water. The mixture was vortexed until fully homogenized and centrifuged at 6,000 g for 5 min. The (6,5) Chirality-enriched bottom phase was collected by aspiration of the top phase and the nanotube interface. Enrichment of the (6,5) chirality was confirmed by UV-vis spectroscopy based on the characteristic E_11_ absorbance peak at 985 nm with minimal contribution from the (6,4) chirality at 880 nm.

### Polymer wrapping of QWNT library

Residual polymers were removed from (6,5)-enriched nanotubes by pressure filtration through a 100 kDa Ultracel disccellulose membrane (Millipore Sigma), with 1% sodium deoxycholate (aq). The procedure was repeated four times to reduce the polymer concentration at least by a factor of 3. To exchange the deoxycholate suspension with the wrapping polymer, 40 µL of polymer solution (10 mg mL^-1^) was added to 200 µL of the nanotube suspension, followed by dropwise addition of 400 µL of methanol under vortexing to precipitate the nanotubes. Subsequently, 40 µL of 5 M sodium chloride was added, and the nanotubes were pelleted by centrifugation at 2,000 g for 10 min. The supernatant was discarded, and the exchange process was repeated once. The final pellet was redispersed in 400 µL of polymer solution (1 mg mL^-1^), and probe-tip sonicated at 30% amplitude for 30 min at 4 °C.^43^ The polymer-wrapped nanotubes were dialyzed against 1X PBS using a 100 kDa dialysis cell (Repligen) to remove unbound polymer by at least a factor of 9. Three ssDNA, (GT)_15_, (AT)_15_, and CT_2_C_3_T_2_C (Integrated DNA Technologies), and two PEG-lipid amphiphiles, DMG-PEG(2000) and DSPE-PEG(2000)-amine (Fisher Scientific), were applied independently to generate 20 polymer-QWNT and five polymer-pristine carbon nanotube samples, comprising a 25-member library for metabolite corona analysis.

### Zeta potential measurements

Aqueous solution of polymer-QWNTs in phosphate-buffered saline with an optical density at (6,5) E_11_ of 1 was carefully transferred into a DTS1070 folded capillary Zetasizer Cell (Malvern, M00049654). Zeta potential values were acquired with the Malvern Zetasizer Nano Z. Instrument parameters for our dispersant are as follows: temperature 25 °C, 0.8872 cP viscosity, refractive index of 1.330, and 78.5 dielectric constant. Equilibrium time is set to 120 seconds. The zeta potentials were calculated with the Smoluchowski model.

### Confocal Raman measurement

Solid-state nanotube samples for Raman analysis were prepared from aqueous dispersions as follows. Briefly, 100 µL of nanotube suspension (OD 1) in an aqueous solution of sodium deoxycholate (1%) was mixed with 1900 µL of ethanol, followed by centrifugation at 13,000 g for 5 min. The supernatant was discarded, and the pellet was deposited onto a glass substrate and dried overnight in a fume hood. Raman spectra were acquired using a QONTOR Confocal Raman spectrometer (Renishaw) under 785 nm excitation with a 1200 lines mm^-1^ grating, 40 mW, 1 s integration time, and a 50x objective.

### Spectroscopic characterization

For the absorption and fluorescence measurements, the nanotubes were dispersed in an aqueous solution of polymers or sodium deoxycholate as described in the earlier paragraph. Absorbance spectra of nanotubes were collected by a V-780 UV-Visible/NIR spectrophotometer (Jasco, Inc.). The optical density at (6,5) E_11_ was adjusted to 0.4 to avoid inner filter effects in the fluorescence measurements. Fluorescence emission spectra of nanotubes were acquired by an InGaAs spectrometer (IMA-SWIR, Photon. etc.). The samples were excited with a 577 nm laser (500 mW, Optoelectronics Tech., Co., Ltd). The light path was shaped and fed into the back of an inverted IX73P2F microscope (Olympus), where it passed through an 850 nm long-pass emission filter, a 20x/0.45 IR objective (Olympus), and illuminated the samples in a 384-well clear flat-bottom UV-transparent microplate (Corning). Following the acquisition, the data were processed using custom MATLAB codes that applied the spectral corrections and background subtraction, and the fluorescence emission peaks were fitted with Lorentzian functions.

### Metabolite corona formation and isolation

Pooled human plasma (IPLAWBNAE100ML, Innovative Research Inc.) was thawed and centrifuged at 13,000 g at 4 °C for 10 min to remove large lipid aggregates. Polymer-wrapped QWNTs (8 µg mL^-1^), pooled human plasma, and 1X PBS were combined in a total volume of 3 mL. The mixture was incubated at room temperature for 2 h at 200 rpm on an orbital shaker. After incubation, the nanotube-plasma mixture was centrifuged at 10,000 g at 4 °C for 25 min, and the supernatant was carefully aspirated. The pellet was resuspended in 1.25 mL of 1X PBS, and the wash and centrifugation steps were repeated three times to remove unbound biomolecules. Pellets not processed immediately were stored at −80 °C until extraction.

### Metabolite extraction from corona pellets

Metabolite extraction was conducted using five extraction solvents (**Table 1**). For single-phase extraction, corona pellets were resuspended in 50 µL of extraction solvent and vortexed for 10 s, followed by bath sonication for 10 min at 20 °C. The suspension was centrifuged at 13,000 g for 10 min at 20 °C. The supernatant was transferred to a 200 µm PTFE membrane filter (Sigma Aldrich), and an additional 100 µL of extraction solvent was loaded onto the filter. The filter was centrifuged at 12,000 g for 10 min, followed by an additional 50 µL of extraction solvent and centrifuged at 12,000 g for 10 min.

For biphasic extraction, corona pellets were resuspended in 500 µL of extraction solvent and shaken at 2,000 rpm for 1 min, then incubated at 18 °C for 10 min. The suspension was centrifuged at 2,500 g for 10 min to separate the top aqueous phase (T) and the bottom organic phase (B). Each phase was carefully transferred into 1.5 mL polypropylene tubes. 250 µL of each phase was transferred to a 200 µm PTFE membrane filter and centrifuged at 12,000 g for 10 min. This step was repeated once with the remaining 250 µL. The combined elutes were dried under vacuum using a SpeedVac concentrator and stored at −80 °C until the LC-MS/MS analysis.

### Addition of Chromatography Internal Standard

For the reverse phase, 50 µL of internal standard, Lipid SPLASH Mix (Cat No: 330707, Avanti Lipids), was added for every 3000 µL of 75% isopropanol. HILIC internal standard is prepared by the Systems Mass Spectrometry core at

Georgia Tech. For HILIC, 50 µL HILIC SPLASH was added for every 3000 µL of 75% methanol. The internal standard-solvent mixtures were vortexed for 30 s. The addition of 75% solvent was to prevent the rapid evaporation of the internal standard over the LC-MS/MS run.

Dried metabolite extracts that were compatible with reverse phase chromatography (RP) were suspended in 100 uL of RP internal standard solution. The metabolite extracts for the hydrophilic interaction column (HILIC) were suspended in 100 µL HILIC internal standard solution.

### LC-MS/MS Analysis

Samples were analyzed by LC-MS/MS using an Exploris 240 Orbitrap mass spectrometer (Thermo Fisher Scientific) for reverse phase (RP) chromatography and a Thermo IQX mass spectrometer (Thermo Fisher Scientific) for HILIC chromatography. Both chromatographic modes were analyzed in positive and negative ionization modes, with scan ranges of 150-2000 *m/z* for RP and 60-900 m/z for HILIC, respectively. Run time per sample is 12 minutes.

For RP chromatography, metabolites were separated on a UPLC C30 column (150 mm × 2.1 mm, 2.7 µm, Thermo Fisher Scientific) at 50 °C with a flow rate of 0.4 mL/min and an injection volume of 2 µL. Mobile phase A consisted 60% acetonitrile, 40% water, 10 mM ammonium formate, and 1% formic acid (positive ion mode) or 10 mM ammonium acetate (negative ion mode). Mobile phase B consisted of 90% isopropanol, 10% acetonitrile, 10 mM ammonium formate, and 1% formic acid (positive ion mode) or 10 mM ammonium acetate (negative ion mode). The gradient was held at 80:20 (A:B) at 0 min, ramped to 0:100 at 8.2 min, and returned to 80:20 over a 12 min total run.

For HILIC chromatography, metabolites were separated on a BEH Z-HILIC (100 mm x 2.1 mm, 1.7 µm, Thermo Fisher Scientific) at 30 °C with an injection volume of 1 µL. Mobile phase C consisted of water, 20 mM ammonium acetate, and adjusted pH 9 with ammonium hydroxide. Mobile phase D consisted of 90% acetonitrile and 10% mobile phase C. The gradient was held at 0:100 (C:D) at 0 min, ramped to 30:70 at 5.5 min, 70:30 at 9 min, and 0:100 at 10 min over 14 min total run.

Metabolite features were subjected to data-dependent acquisition, and MS/MS spectra were matched to a reference spectral library for putative metabolite annotations.

### LC-MS/MS data processing and feature selection

LC-MS raw data were processed in Compound Discoverer 3.3 (Thermo Fisher Scientific) separately for each chromatographic mode (RP and HILIC) and ionization polarity (positive and negative). Feature detection, alignment, and spectral library matching were performed using default Compound Discoverer workflows. Feature lists from HILIC and RP modes were curated by parsing m/z values and removing duplicate entries with a mass tolerance of 0.02 Da and a retention time tolerance of 0.3 min, retaining the feature with the highest mean intensity.

Gap status was assigned per feature per sample: “no gap” (detected above the signal threshold), “missing ion” (partially detected across replicates), or “full gap” (completely absent from all replicates in a sample group). Features classified as full gap across all three technical replicates within a sample group were discarded, as the complete absence of signal precludes distinguishing true biological absence from technical dropout. Missing values in the remaining features were imputed using half the minimum observed intensity for that feature across all non-missing samples, followed by row-median normalization to correct for inter-sample loading variability.

A tiered statistical filtering strategy was then employed to maximize the retention of biological information while maintaining high reproducibility. Tier 1 retained features exhibiting a coefficient of variation (CoV) < 30% across technical triplicates. Tier 2 retained features with higher variance (CoV ≥ 30%) only if they demonstrated a pronounced biological signal, defined as |log_2_FC| > 1.0 relative to the plasma reference control. This tiered approach removed low-intensity stochastic noise while preserving high-variance but biologically significant markers. The final metabolomics library was generated by combining unique survivors from both HILIC and RP modes across all four analytical configurations (RP±, HILIC±).

### Inverse-variance weighted log_2_ fold change estimation

Metabolite features were quantified across five independent extraction methods (Table 1), each producing distinct systematic offsets in absolute ion intensities. To combine these measurements without confounding by between-method batch effects, we employed an inverse-variance weighted (IVW) meta-analysis analogous to fixed-effect meta-analysis in clinical epidemiology.

For each feature i and nanotube condition j, the log_2_ fold change relative to plasma was computed independently within each extraction method k:

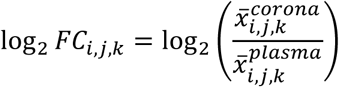

where *x̅* denotes the group mean intensity. The associated standard error SE_i,j,k_ was estimated from the within-group variances and sample sizes of the corona and plasma groups for method k, allowing for unequal variance (Welch approximation).

The IVW-combined estimate across K extraction methods was then computed as follows:

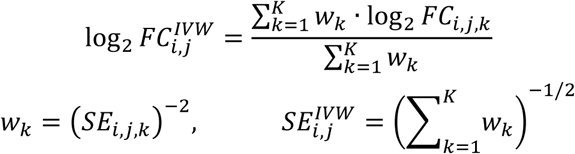

The combined p-value was obtained from a two-tailed z-test.

This procedure was applied to assess two experimental contrasts. First, for each of the 25 nanotube types, the corona-versus-plasma IVW log_2_FC quantifies the overall enrichment or depletion of each feature in the nanotube corona. Second, for the QWNT-versus-pristine comparison, the log_2_FC was computed as the difference between each QWNT and the pristine baseline within the same polymer wrapping, isolating the defect-specific modulation of corona composition from the polymer effect. Features with fewer than two extraction methods contributing valid measurements were excluded. No multiple-testing correction was applied at this stage, and false discovery rate (FDR) correction was applied downstream within each enrichment analysis independently.

To obtain a single enrichment score per feature across all nanotube types, the 25 per-condition IVW log_2_FC values were averaged using an arithmetic mean. Significance of the consensus log_2_FC was assessed by a one-sample t-test against zero (two-sided), using the observed variance across the 25 conditions as the error term. For features detected in only one condition (n = 1), significance was estimated by an IVW z-test using the propagated standard error from the first-stage combination. Features with no valid log_2_FC in any of the 25 coronas were excluded from the consensus estimate. This metric is used for volcano plots of **Fig 4c,d** and class-level enrichment analyses of **Fig 4f,g**. All polymer-specific and QWD-specific analyses use the individual per-condition IVW values directly.

### Batch-To-Batch Reproducibility Assessment

Two independent experimental batches of (AT)_15_ wrapped, *N,N*-diethyl-4-aminoaryl functionalized carbon nanotubes were used. For each batch, peak areas were first normalized by the mean plasma intensity per extraction method per run, then median-normalized across matched metabolite features (m/z ±0.02 Da, RT ±0.3 min), and full-gap features were removed. The inter-batch coefficient of variation was computed from normalized intensities across batches. Principal component analysis was performed on log_2_-transformed intensities (normalized to the global feature median) to confirm that samples are clustered by analytical batch. Relative log expression plots were generated from log_2_-transformed intensities to detect systematic intensity shifts between batches.

### Machine Learning Classification of Metabolite Features

Spectral library annotation in Compound Discoverer assigned metabolite class labels to 677 RP features and 1,081 HILIC features prior to deduplication. To extend class annotations to the remaining unannotated features, supervised machine learning classifiers were trained using three instrument-level descriptors: mass-to-charge ratio (m/z), mass defect, and ESI polarity. Metabolite classes were consolidated into 5 classes for RP (phospholipids, complex lipids, lysophospholipids, acylcarnitines, fatty acids) and 9 classes for HILIC (phospholipids, organic acids, fatty acids & amides, secondary & xenobiotic, amino acids & peptides, lysophospholipids, acylcarnitines, complex lipids, benzenoids) based on chromatographic retention behavior, mass spectral properties, and biosynthetic relationships (Supplementary Table 4).

Three classifiers, k-nearest neighbors (KNN), random forest (RF), and gradient boosting (XGBoost), were trained using 3-fold cross-validation on 80% of annotated features and evaluated on a held-out 20% test set, stratified by class. For RP, the RF model achieved a weighted F1 score of 0.934. For HILIC, the best-performing RF model achieved a weighted F1 of 0.525, reflecting greater spectral overlap among polar metabolite classes. Features predicted with confidence below 0.40 were flagged as Low Confidence.

To account for differences in classification reliability across HILIC classes, a tiered analysis framework was implemented. Track 1 classes (per-class F1 ≥ 0.56 and fraction of dataset > 5%), Phospholipids, Lysophospholipids, Organic Acids, Fatty Acids & Amides, and Secondary & Xenobiotic, were analyzed using combined annotated and predicted features. Track 2 classes, including Acylcarnitines, Complex Lipids, Benzenoids, and Amino Acids & Peptides, were analyzed with annotated and predicted features reported separately and compared for consistency. All RP classes were treated as Track 1.

### Feature Deduplication

Untargeted LC-MS frequently detects the same metabolite as multiple features corresponding to different adducts (e.g., [M+H]⁺, [M+Na]⁺, [M+K]⁺) or in-source fragments with near-identical m/z and retention time. To prevent such redundant entries from inflating class-level statistics and pathway enrichment counts, a name-based deduplication step was applied after ML classification but before all class-level analyses.

Deduplication rules were as follows. Annotated features sharing a common base metabolite name (after stripping trailing numeric suffixes such as “.1”, “.2”) were assigned to the same deduplication group. Unannotated features (assigned numeric identifiers by Compound Discoverer) were never collapsed, as their chemical identity cannot be confirmed. Features with generic prefix annotations (e.g., “PC.4”, “FA.11”) were also excluded from collapsing to avoid incorrectly merging structurally distinct species. Within each deduplication group, a single representative feature was retained by computing the median IVW log₂FC across group members; p-values were taken as the minimum (most significant) across members. Deduplication reduced the feature count to 6,383 (RP, from 6,453; 1.1% reduction) and 2,529 (HILIC, from 2,732; 7.4% reduction), with acylcarnitines and phospholipids most affected due to the prevalence of sodium and potassium adducts in these classes. ML classification and IVW fold-change estimation were performed on the full feature set prior to deduplication.

### Class-Level Enrichment Analysis

For each metabolite class and nanotube condition, the distribution of per-feature IVW log_2_FC values was summarized by the median and interquartile range (Q25–Q75). Per-feature significance was determined by the IVW p-value (two-sided z-test, as described above). At the class level, the statistical significance of the median enrichment was assessed using the Wilcoxon signed-rank test against zero, with p-values corrected for multiple testing using the Benjamini–Hochberg (BH) procedure for each comparison type (corona vs plasma or QWD vs pristine). Significance thresholds were set at q < 0.05.

For the QWD-versus-pristine comparison, the per-feature median Δlog_2_FC across all 20 QWD conditions (5 polymers × 4 defects) was first computed, followed by class-level aggregation. Class sensitivity to QWDs was quantified as the fraction of features within each class that were significantly affected (p < 0.05, |Δlog_2_FC| > 0.5) by at least one of the 20 polymer-QWNT combinations.

### Enrichment Score Analysis

To assess whether Track 2 HILIC classes were non-randomly distributed within ranked feature lists — without relying on classification accuracy for individual class membership — a pre-ranked enrichment analysis was performed. For each nanotube condition, all HILIC features were ranked by their IVW-combined log_2_FC (descending). A running enrichment score was computed by walking down the ranked list: at each class member, the score incremented proportionally to |log_2_FC| (weighted hit), and at each non-member, the score decremented uniformly (miss penalty). The enrichment score (ES) was defined as the maximum absolute deviation from zero, with positive ES indicating that class members are concentrated at the corona-enriched end of the ranked list and negative ES indicating that they are concentrated at the plasma-enriched end. This procedure was applied per polymer condition (for polymer-effect ES analysis, Fig 5f) and per defect type (for QWD-effect ES analysis, Supplementary Fig 25), using the mean log_2_FC across all five polymers for each defect.

### DNA versus PEG Comparison

To quantify the effect of polymer wrapping type on corona composition, a pairwise comparison was performed between DNA-wrapped and PEG-wrapped pristine nanotubes. For each feature, the log_2_FC difference was computed as the average of the three DNA pristine conditions minus the average of the two PEG pristine conditions. Statistical significance was assessed by Welch’s t-test when both groups had ≥ 2 non-missing values; for features with fewer observations, a z-test based on IVW-propagated standard errors was used. No multiple-testing correction was applied at the feature level; class-level summaries were interpreted based on effect-size consistency across features rather than individual feature significance.

### Mummichog Pathway Enrichment Analysis

To characterize the adsorbed metabolome at the pathway level, mummichog (v2.7.0) was applied to all metabolite features detected from any nanotube coronas vs plasma. For each feature in the deduplicated HILIC dataset, the IVW-combined log_2_FC across all 25 nanotube conditions was pooled using the minimum p-value across the five polymers as the significance statistic. Features were split by ESI polarity and ran separately, as mummichog requires ionization mode specification for accurate adduct matching. Input files contained four columns (m/z, retention time, p-value, t-score) with p < 0.05 as the significance cutoff. Mummichog matched m/z values to the human metabolic model (KEGG/BioCyc) with 100 permutations for background estimation. Results from positive and negative modes were combined, retaining pathways with permutation p < 0.05 in either mode.

### Sensitivity Analyses

A deduplication sensitivity analysis was performed by running all class-level enrichment analyses on both the full and deduplicated feature sets. For each class, the direction (enriched vs depleted), magnitude (median log_2_FC), and statistical significance (q-value) were compared between the two analyses. All class-level conclusions were preserved after deduplication (Supplementary Table 6).

### Software and Code Availability

All data processing and statistical analyses were performed in Python 3.12 using pandas (v2.0), NumPy (v1.24), SciPy (v1.11), scikit-learn (v1.3), and XGBoost (v2.0). Figures were generated using GraphPad Prism (v11.0.0) and Biorender. Mummichog (v2.7.0) was used for pathway enrichment analysis. Custom analysis scripts are available at [repository URL/To be shared].

## Author contributions

**M.K.** conceived the project. **J.G.** and **I.H.** conducted the experiments. **J.G.**, **I.H.**, and **M.K.** analyzed data for the article, wrote, reviewed, and edited the manuscript before submission. All authors have given approval to the final version of the manuscript.

## Declaration of Competing Interest

M.K. is a co-founder with equity interest in Nine Diagnostics. Other co-authors declare no competing interests.

## Acknowledgments

This work was supported in part by the National Institutes of Health (R00-EB033580) to M.K. This research was funded, in part, by the Advanced Research Projects Agency for Health (ARPA-H) under Agreement No. 1AY2AX000080-01. The views and conclusions contained in this document are those of the authors and should not be interpreted as representing the official policies, either expressed or implied, of the U.S. Government. J.G. was supported in part by the NSF GAANN Fellowship. I.H. was supported in part by the Sejong Science Fellowship Program (RS-2024-00358020) of the National Research Foundation of Korea. The metabolomics data were acquired at the Systems Mass Spectrometry Core Facility at the Georgia Institute of Technology, funded by the NIH (1S10OD038327-01). Raman data were acquired at the Materials Characterization Facility of the Institute for Matter and Systems at the Georgia Institute of Technology.

