## Supplementary Information for "Chemically Engineered Carbon Nanotubes Map Class-Selective Metabolite Enrichment from Human Plasma"

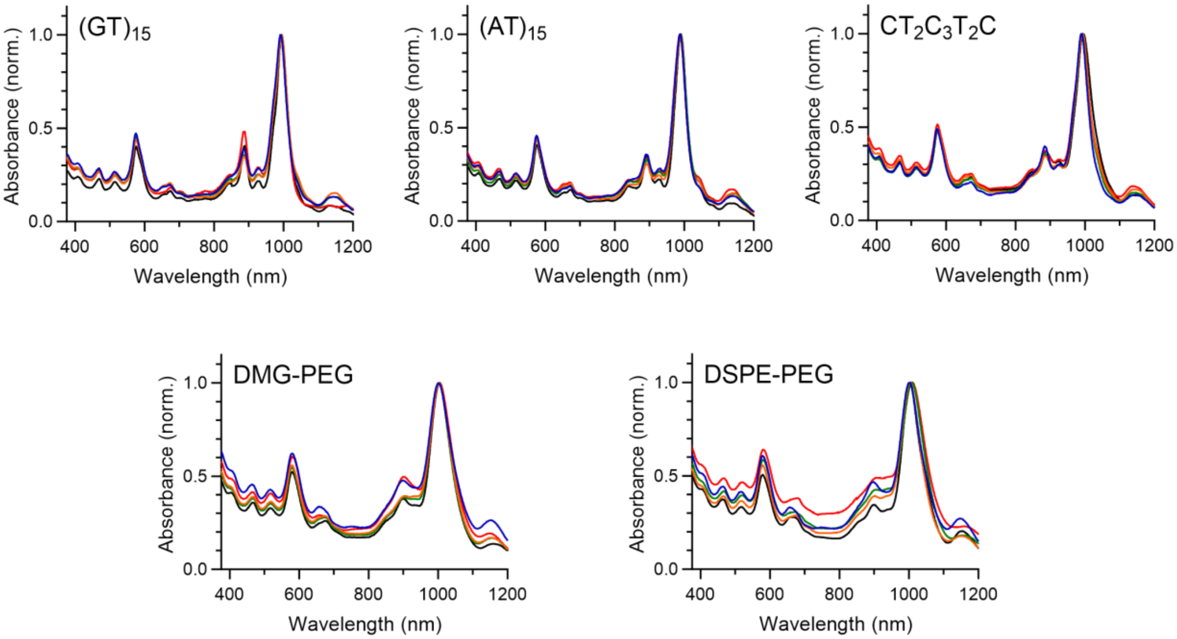

**Supplementary Figure 1.** UV-visible-NIR absorption spectra of polymer-wrapped, (6,5)-enriched carbon nanotubes. Red, orange, green, blue, and black spectra indicate 4-carboxylaryl QWNTs (Carboxyl), N,N-diethyl-4-aminoaryl QWNTs (Amino), 3,5-dinitroaryl QWNTs (Dinitro), 3,4,5-trifluoroaryl QWNTs (Trifluoro), and pristine nanotubes in 1X phosphate-buffered saline, respectively. The absorption spectra are normalized to (6,5) $E_{11}$ absorption band.

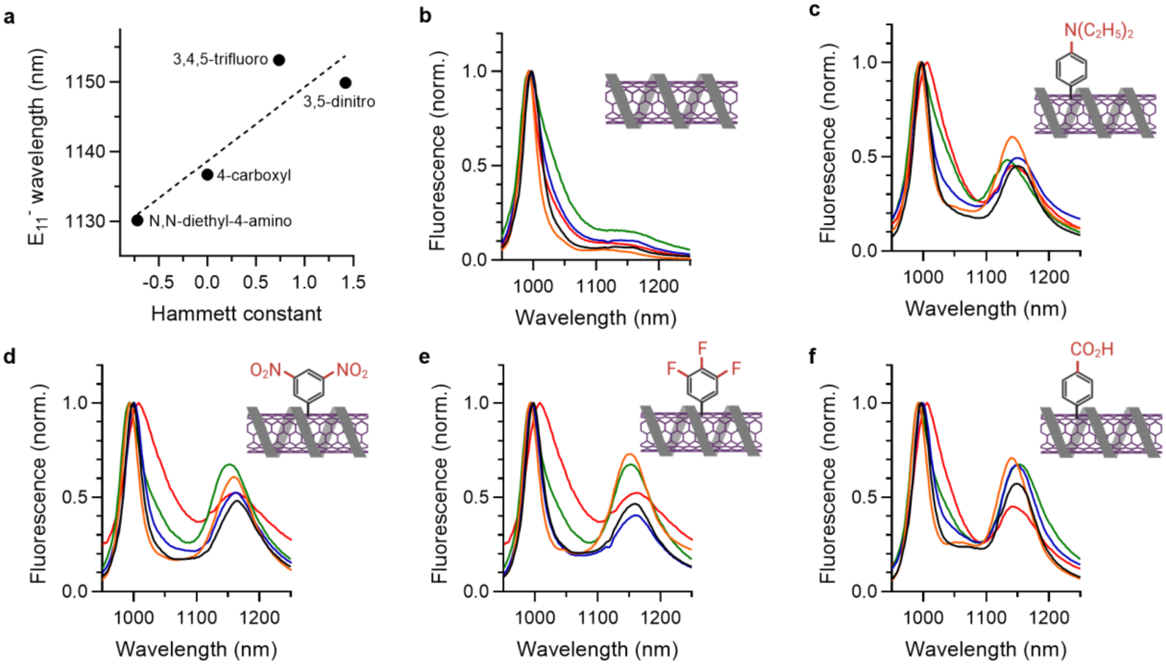

**Supplementary Figure 2.** Fluorescence characterization of (6,5) enriched carbon nanotubes. **a,** Correlation between Hammett constant of aryl QWDs and $E_{11}^{-}$ emission wavelength. Normalized NIR fluorescence spectra of polymer-wrapped **b,** pristine nanotubes and **c-f,** QWNTs. **c,** *N,N*-diethyl-4-aminaryl QWNTs. **d,** 3,5-dinitroaryl QWNTs. **e,** 3,4,5-trifluoroaryl QWNTs. **f,** 4-carboxylaryl QWNTs. Color indicates different polymer wrapping (Blue: $CT_2C_3T_2C$; Black: $(GT)_{15}$; Orange: $(AT)_{15}$; Green: DMG-PEG(2000); Red: DSPE-PEG(2000)-amine). Excitation wavelength is 577 nm.

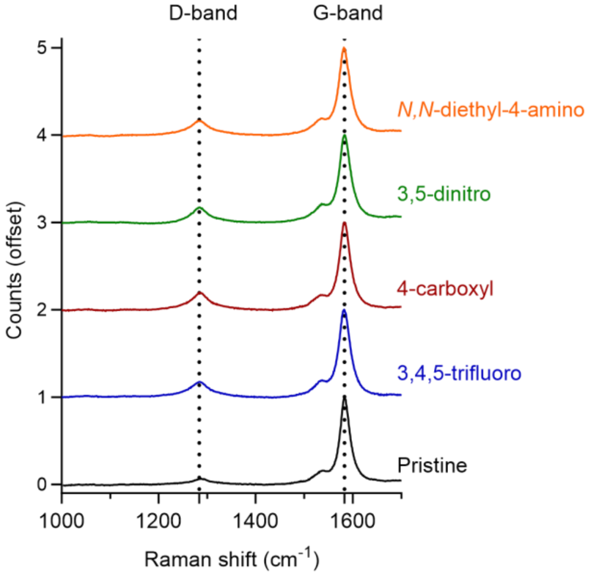

**Supplementary Figure 3.** Raman spectra of pristine and QWD-functionalized nanotubes. Full Dashed lines indicate Raman D and G bands. Each spectrum was acquired with 40 mW of 785 nm laser excitation and normalized with respect to the G-band.

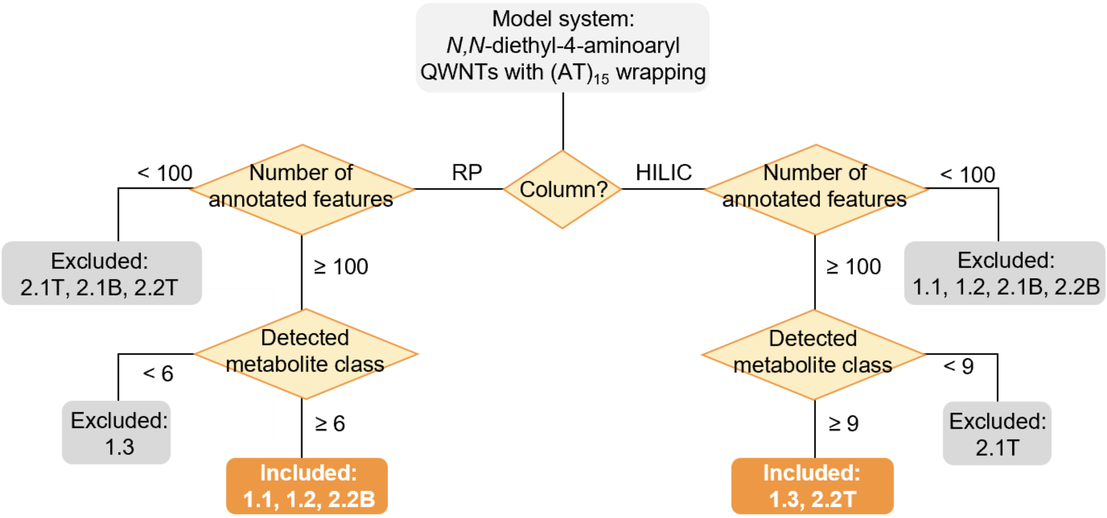

**Supplementary Figure 4.** Extraction methods decision flow. Sequential two-criterion framework applied to a model system, $(AT)_{15}$ wrapped *N,N*-diethyl-4-aminoaryl QWNTs. Methods were retained if they yielded ≥100 annotated features and covered at least 6 (RP) or 9 (HILIC) distinct metabolite classes. Methods that do not meet both criteria are shown in gray.

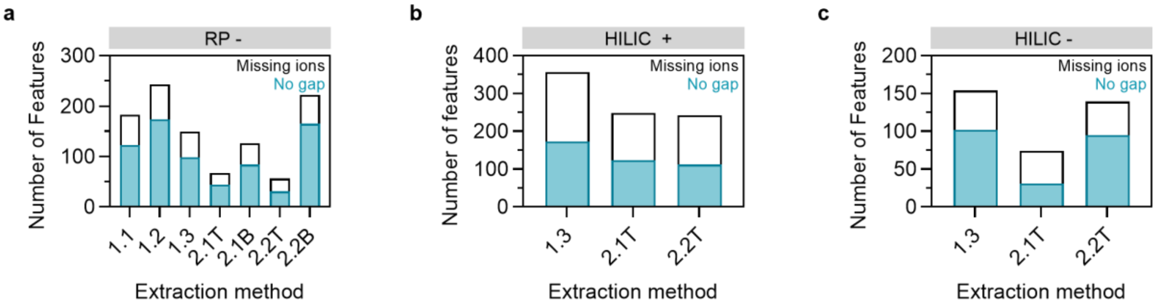

**Supplementary Figure 5.** Feature counts of no gap (blue) and missing ions (white) in selected extraction methods for **a,** RP-, **b,** HILIC+, and **c,** HILIC- mode in $(AT)_{15}$ wrapped, *N,N*-diethyl-4-aminoaryl QWNTs.

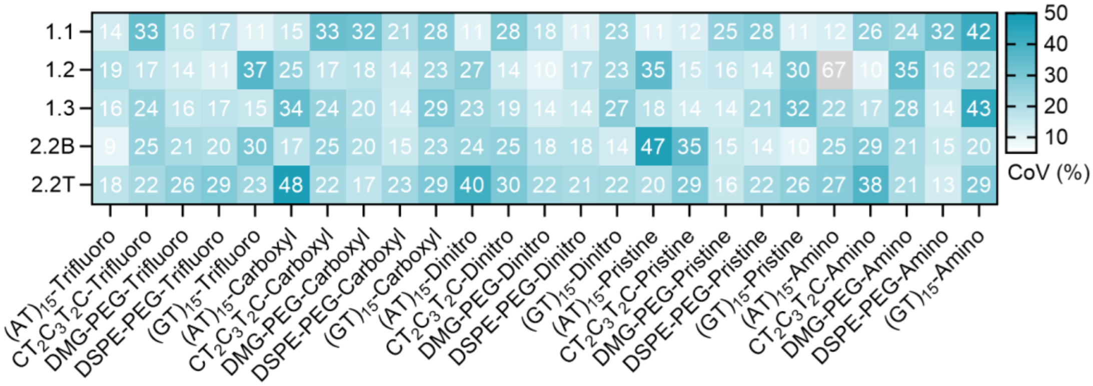

**Supplementary Figure 6.** Intra-batch analytical reproducibility across all nanotube coronas. Coefficient of variation (CoV, %) heatmap for nanotubes and extraction methods.

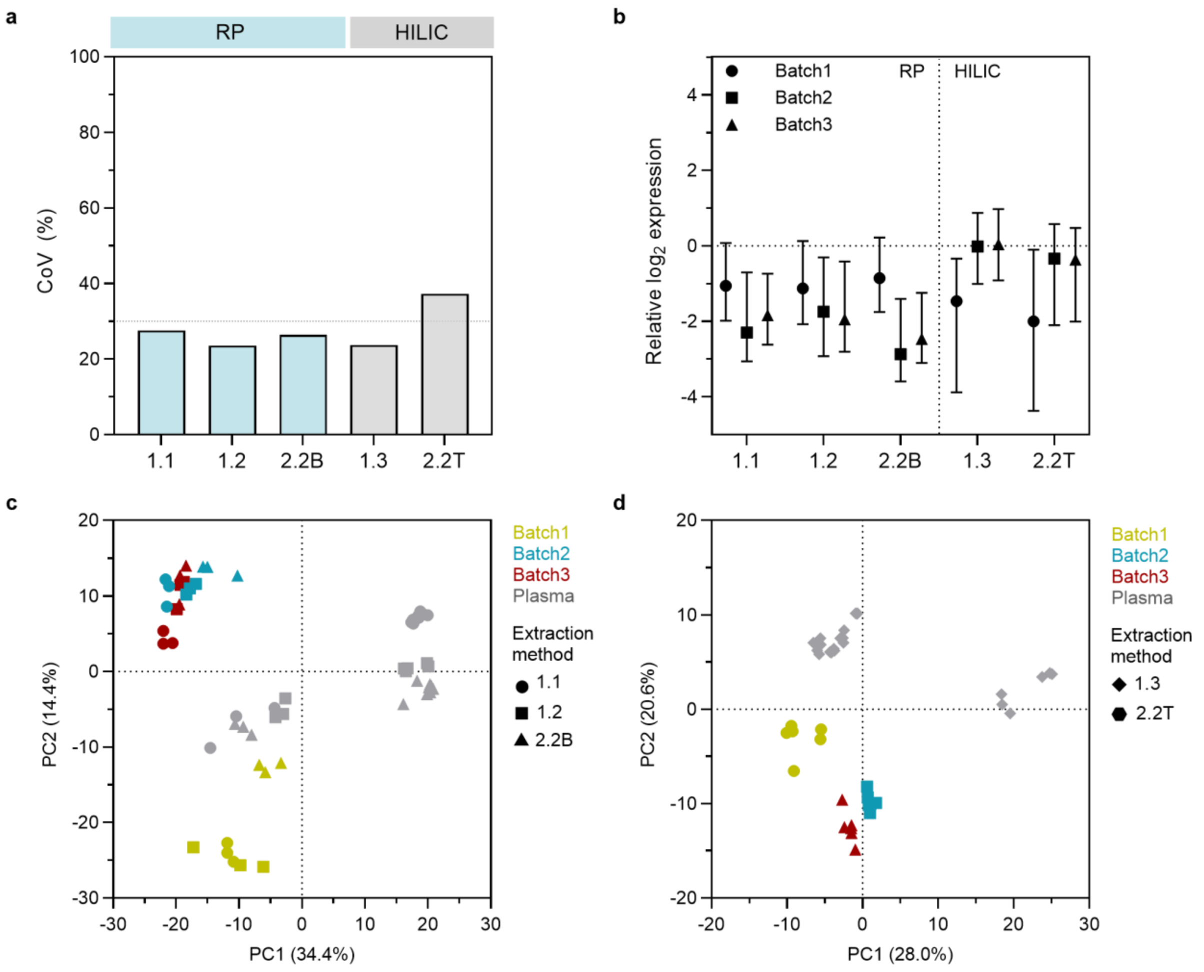

**Supplementary Figure 7.** Inter-batch analytical reproducibility of $(AT)_{15}$-wrapped, *N,N*-diethylaminoaryl functionalized QWNTs across extraction methods in three independently synthesized QWNT batches. **a,** Inter-batch CoV from plasma-normalized intensities across three batches. Dashed line is 30% threshold. **b,** Normalized relative $\log_2$ expression (RLE) analysis per batch and extraction method. Point and error bar denote the median RLE and interquartile range, respectively. Principal component analysis of $\log_2$ intensities for the QWNTs with plasma in **c,** RP, and **d,** HILIC. PCA separation between batches reflects differences in the subset of features detected above the signal threshold in each run, rather than systematic technical bias.

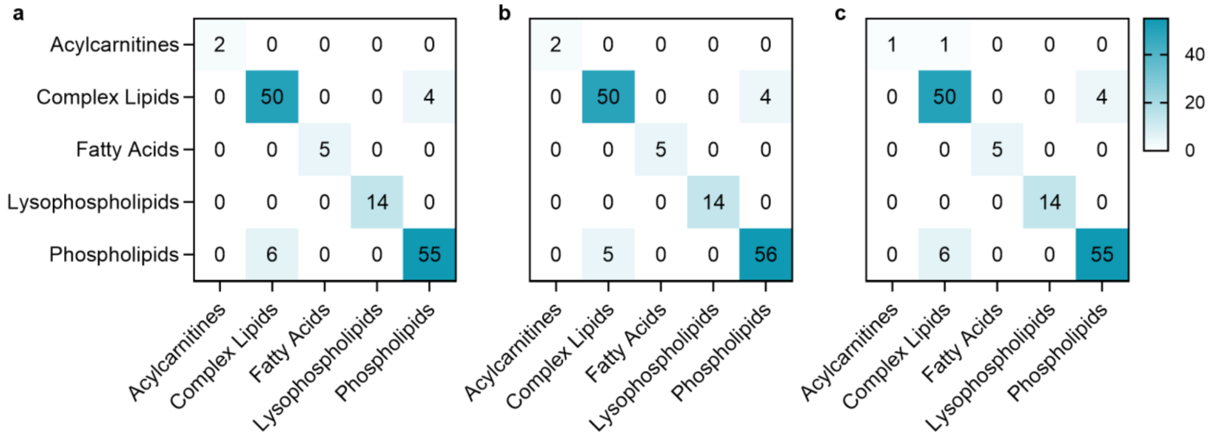

**Supplementary Figure 8.** Confusion matrices for machine learning classifiers on RP-detected metabolites. **a,** XGBoost, **b,** Random forest, and **c,** KNN.

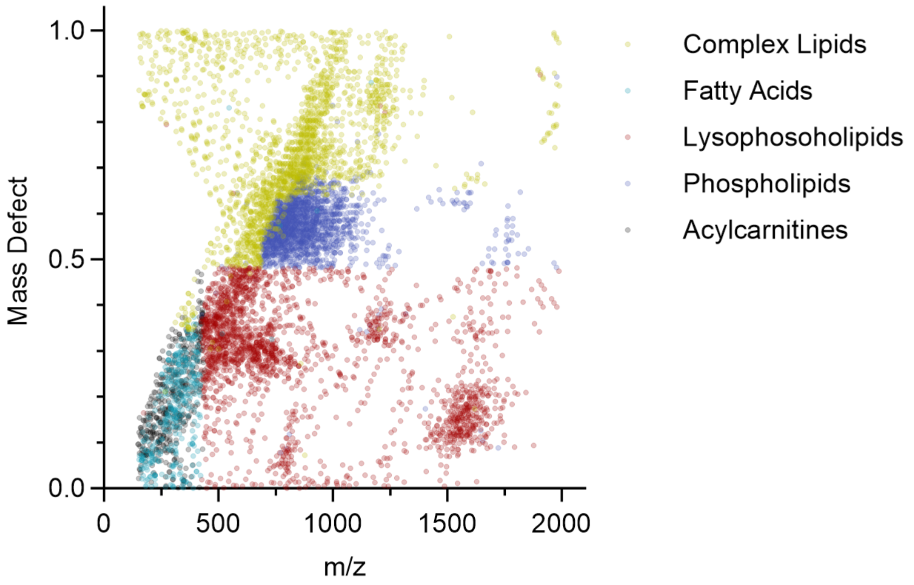

**Supplementary Figure 9.** Feature space distribution of RP metabolites. Scatter plots of m/z versus mass defect for annotated RP features, colored by metabolite class. Complex lipids and phospholipids occupy distinct high m/z ranges, acylcarnitines feature low mass defects, and lysophospholipids are separable from phospholipids by mass defect at overlapping m/z values.

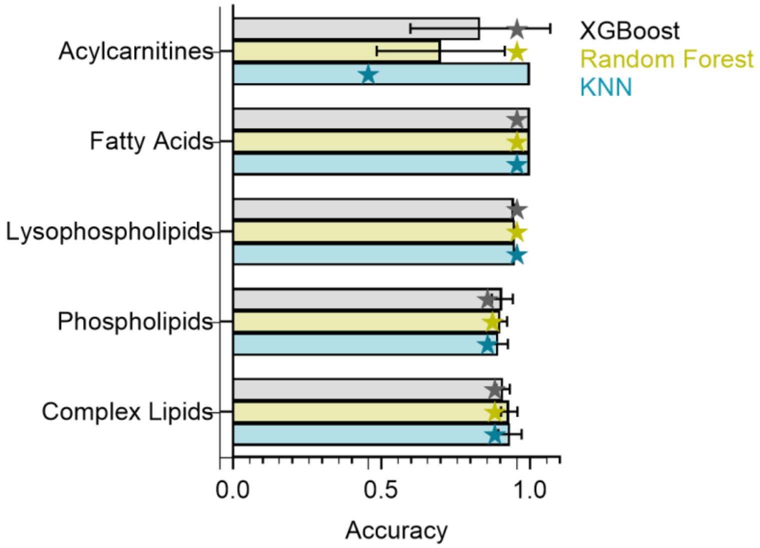

**Supplementary Figure 10.** Per-class classification accuracy for RP mode in the optimized multi-class classifier. Error bars indicate the standard deviation of 3-fold cross-validation. Star symbols are the test set score.

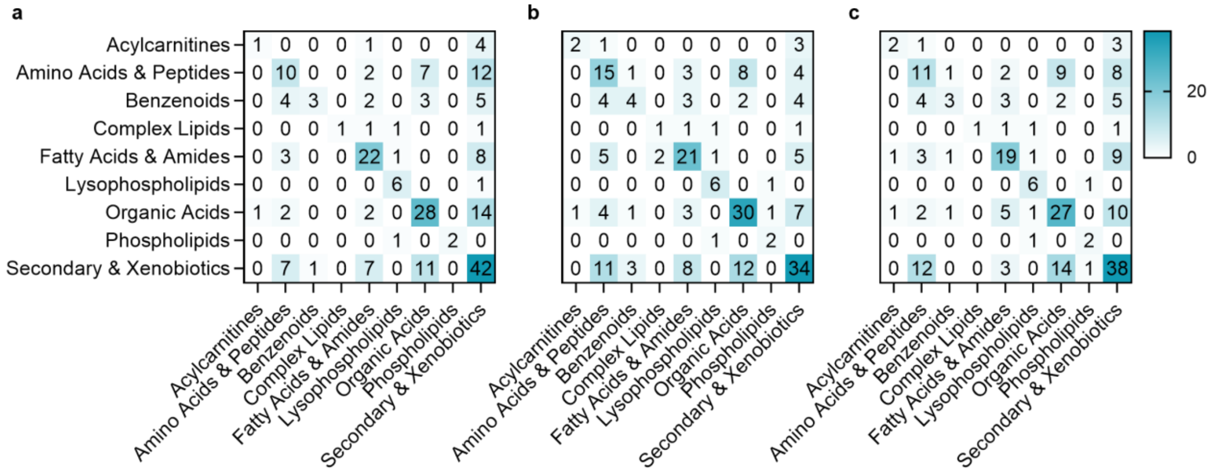

**Supplementary Figure 11.** Confusion matrices for machine learning classifiers on HILIC-detected metabolites. **a,** XGBoost, **b,** Random forest, and **c,** KNN.

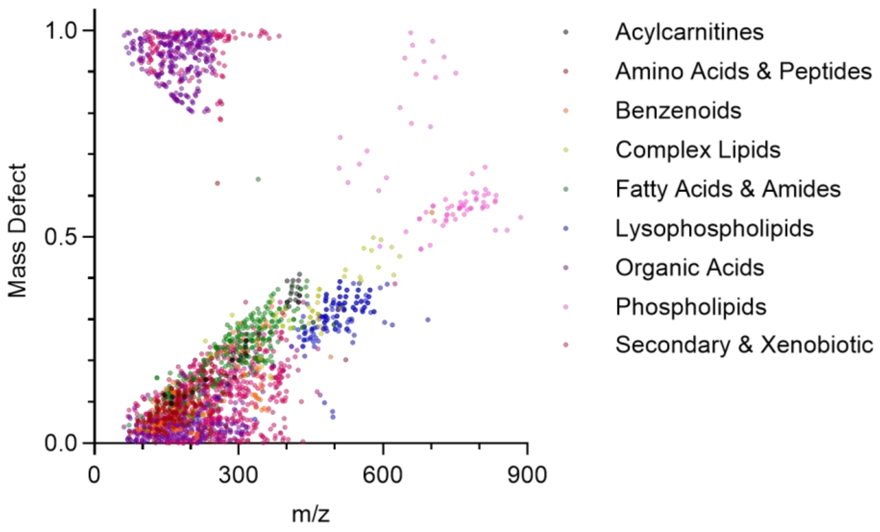

**Supplementary Figure 12.** Feature space distribution of HILIC metabolites. Scatter plots of m/z versus mass defect for annotated HILIC features, colored by metabolite class.

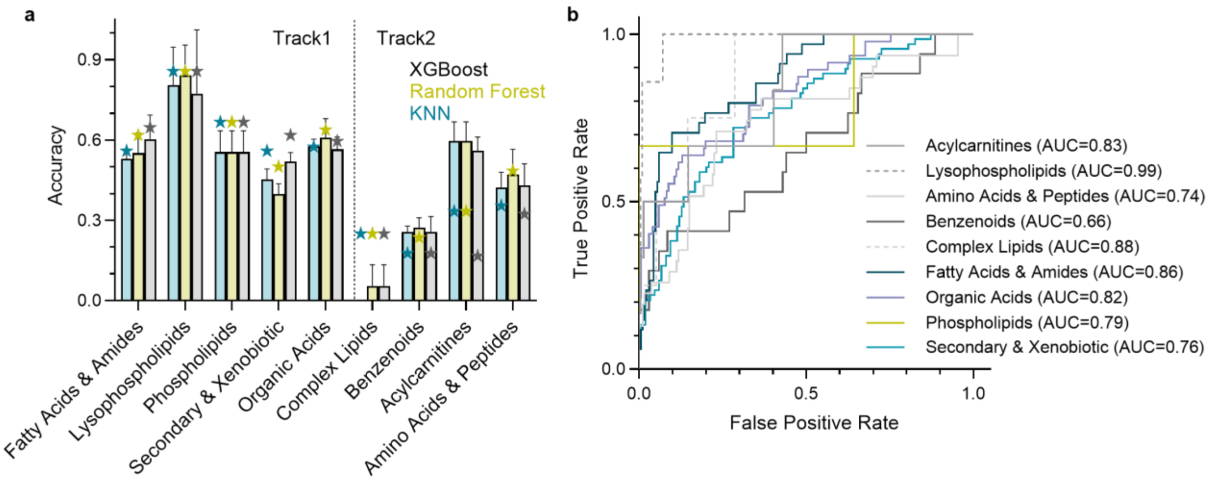

**Supplementary Figure 13.** Per-class classification performance for HILIC mode. **a,** Per-class classification accuracy in the optimized multi-class classifier. Error bars indicate the standard deviation of 3-fold cross-validation. Star symbols are the test set score. **b,** per-class test ROC curves for the best-performing random forest classifier.

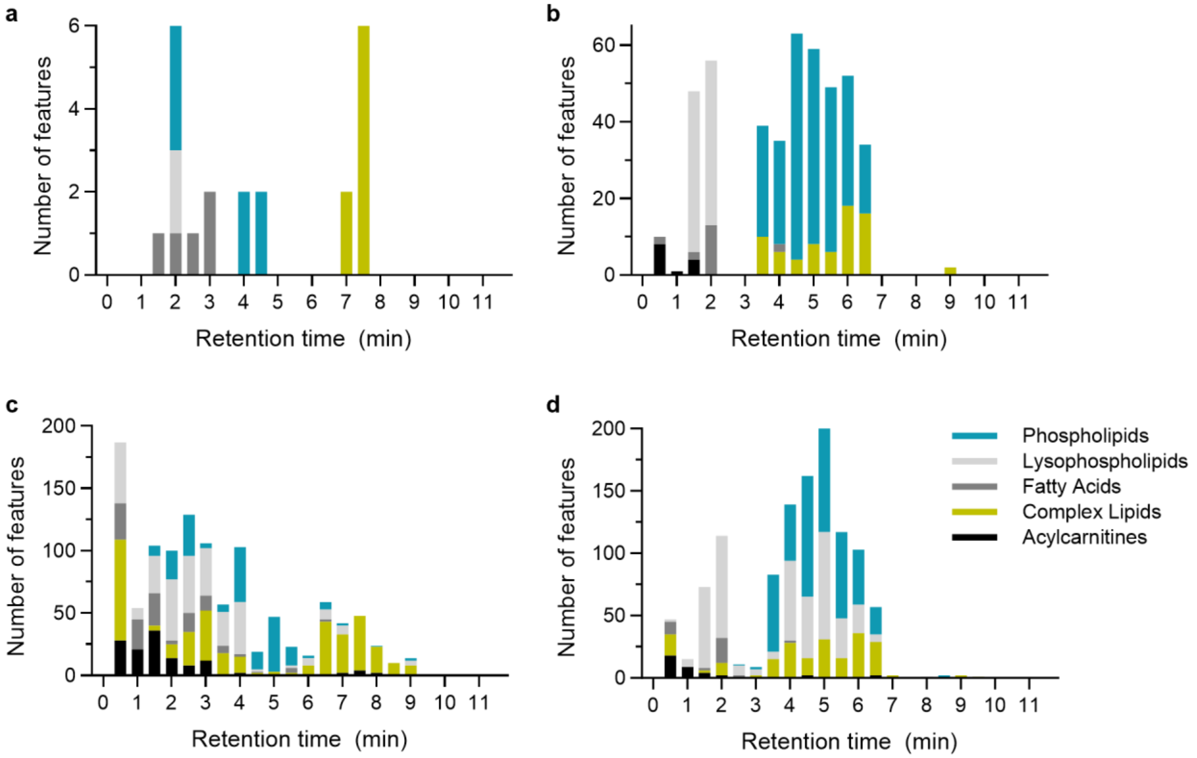

**Supplementary Figure 14.** Retention time distributions of significantly enriched features ($p < 0.05$, $|\log_2 FC| > 1$) for RP mode. Annotated features in **a,** corona, and **b,** plasma. Annotated and predicted features in **c,** corona, and **d,** plasma. Colors indicate metabolite class.

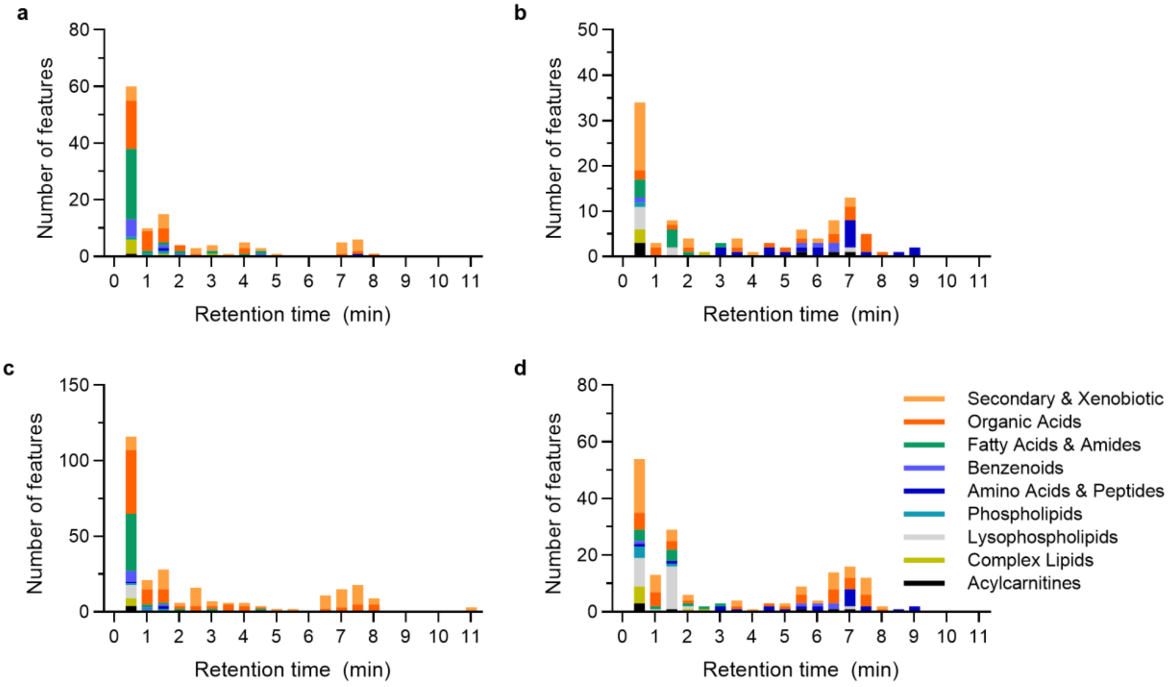

**Supplementary Figure** 15**.** Retention time distributions of significantly enriched features ($p < 0.05$, $|\log_2 FC| > 1$) for HILIC mode. Annotated features in **a,** corona, and **b,** plasma. Annotated and predicted features in **c,** corona, and **d,** plasma. Colors indicate metabolite class.

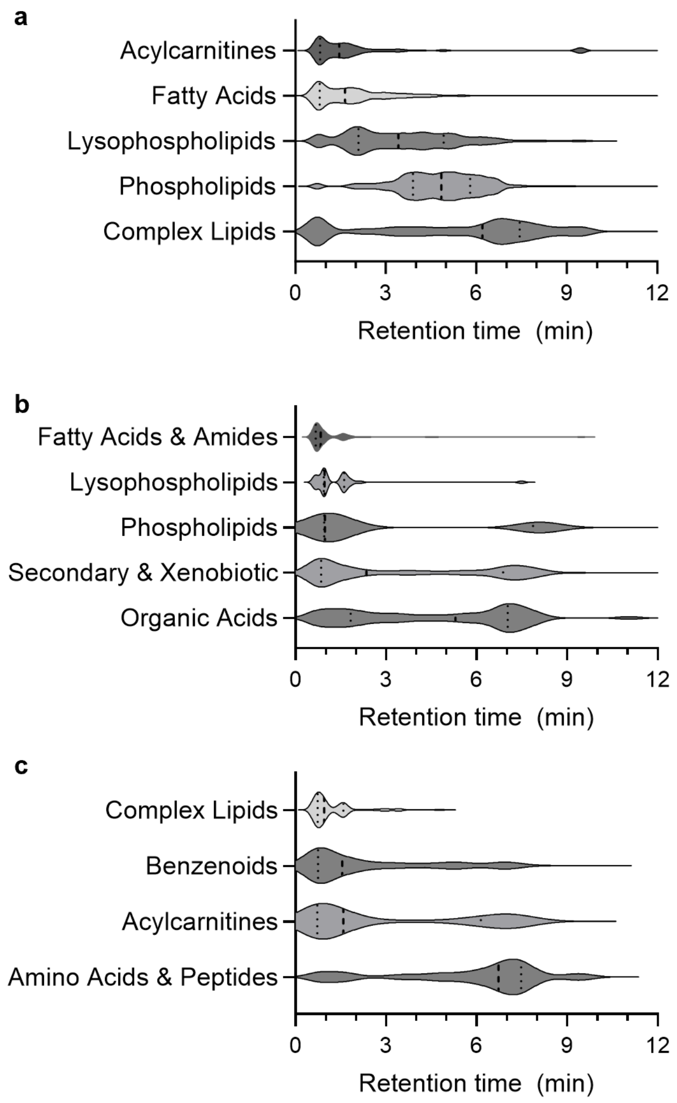

**Supplementary Figure 16.** Violin plots of median retention time distribution of Track 1 classes in **a,** RP and **b,** HILIC and **c,** HILIC Track 2 classes. The distribution confirms the differential enrichment of nonpolar vs. polar analytes across all retention time profiles. Dashed and dotted vertical lines are median and interquartile range, respectively.

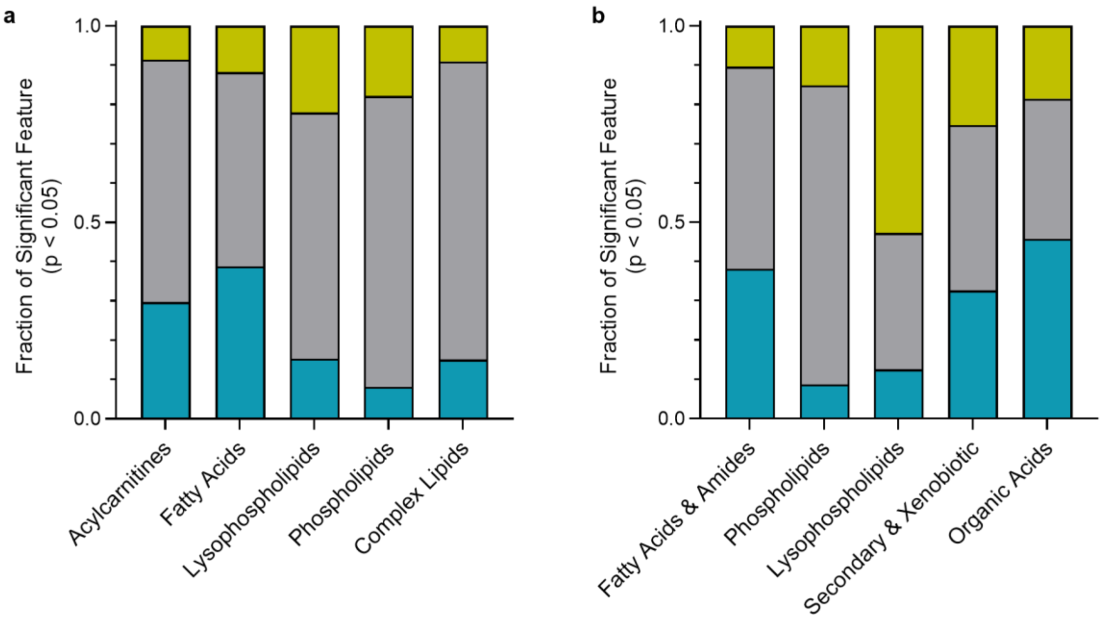

**Supplementary Figure 17.** Per-class significance proportions in the corona versus plasma comparison. Stacked bar plots showing the fraction of features within each metabolite class that are significantly corona-enriched (blue), plasma-enriched (yellow), or not significant (gray) at $p < 0.05$. **a,** RP. **b,** HILIC Track 1.

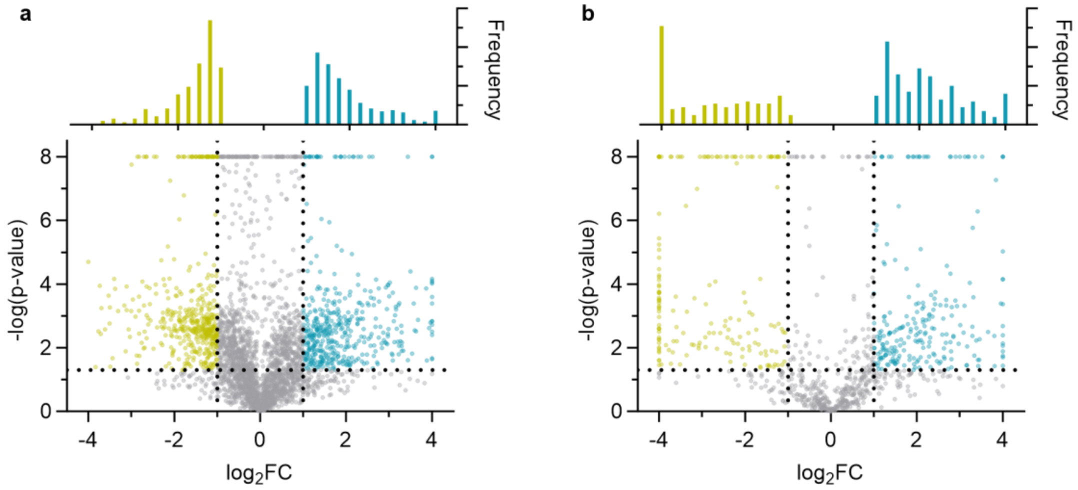

**Supplementary Figure 18.** Volcano plots comparing pristine nanotube corona to plasma for **a,** RP and **b,** HILIC. $Log_2FC$ represents pristine nanotube corona enrichment minus plasma enrichment. Blue and yellow points are corona-enriched and plasma-enriched features, respectively, that exceed the significance thresholds (p-value cutoff of 0.05 and $|log_2FC|$ cutoff of 1, dashed lines). The top panels are the frequency distribution of significant features.

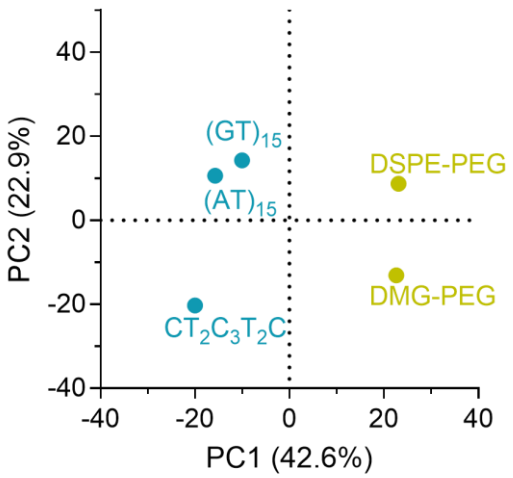

**Supplementary Figure 19.** Principal component analysis of HILIC corona profiles (Track 1) across pristine polymer wrappings.

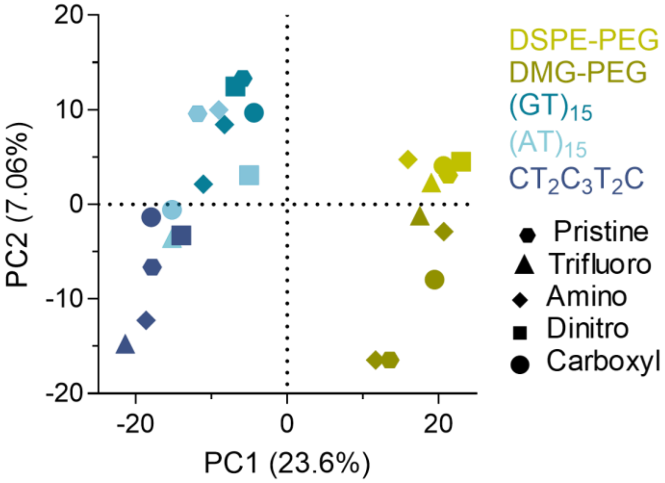

**Supplementary Figure 20.** Principal component analysis of HILIC corona profiles (Track 1) across all 25 nanotube conditions (20 polymer-QWNTs + 5 pristine). Marker shape and color indicate QWD type and polymer type, respectively.

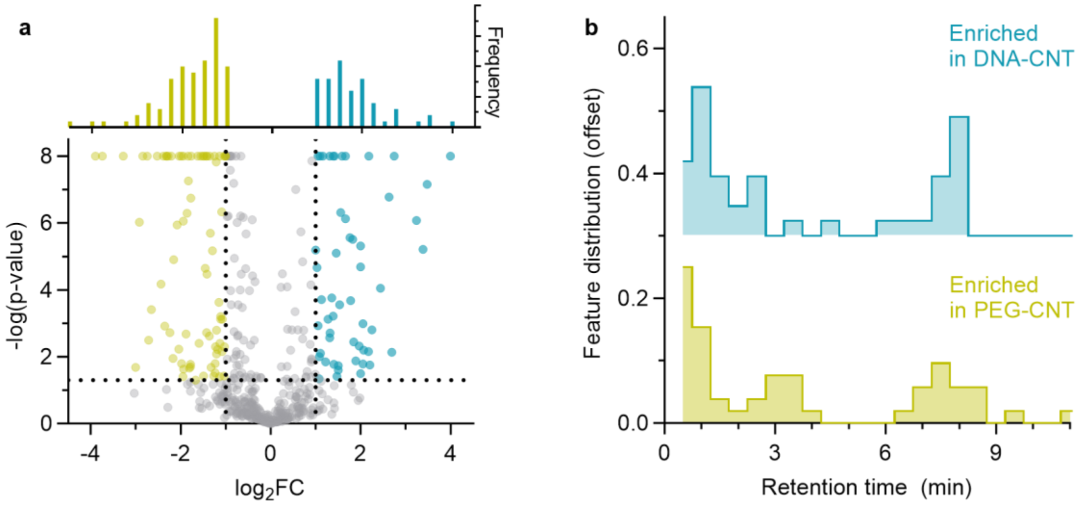

**Supplementary Figure 20.** Polymer class-dependent metabolite enrichment with pristine nanotubes in HILIC. **a,** Volcano plot comparing coronas extracted from DNA-wrapped to PEG-wrapped pristine nanotubes. $Log_2FC$ represents IVW-DNA corona enrichment minus IVW-PEG corona enrichment. Blue and yellow points represent features significantly enriched in DNA-CNT and PEG-CNT coronas, respectively ($p < 0.05$, $|log_2FC| > 1$). The top panel is the frequency distributions of significant features. **b,** Retention time distribution of significant features detected in **a**.

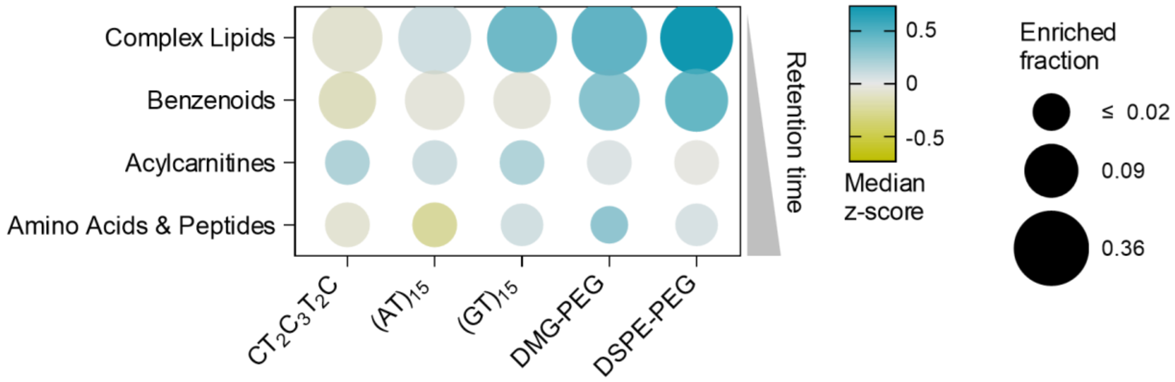

**Supplementary Figure 21.** Class-level enrichment bubble plot of annotated features in HILIC Track 2. Dot size indicates the enriched fraction, and color indicates the median z-score of $log_2FC$ (corona − plasma) normalized across polymers within each class.

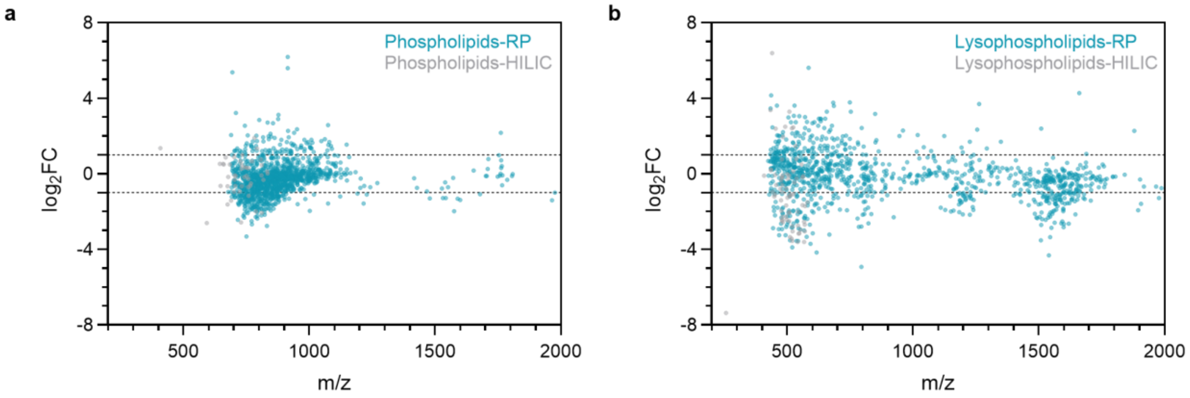

**Supplementary Figure 22.** m/z distributions of **a,** phospholipid and **b,** Lysophospholipid features in RP and HILIC modes. $\log_2$FC represents the difference between corona enrichment and plasma enrichment.

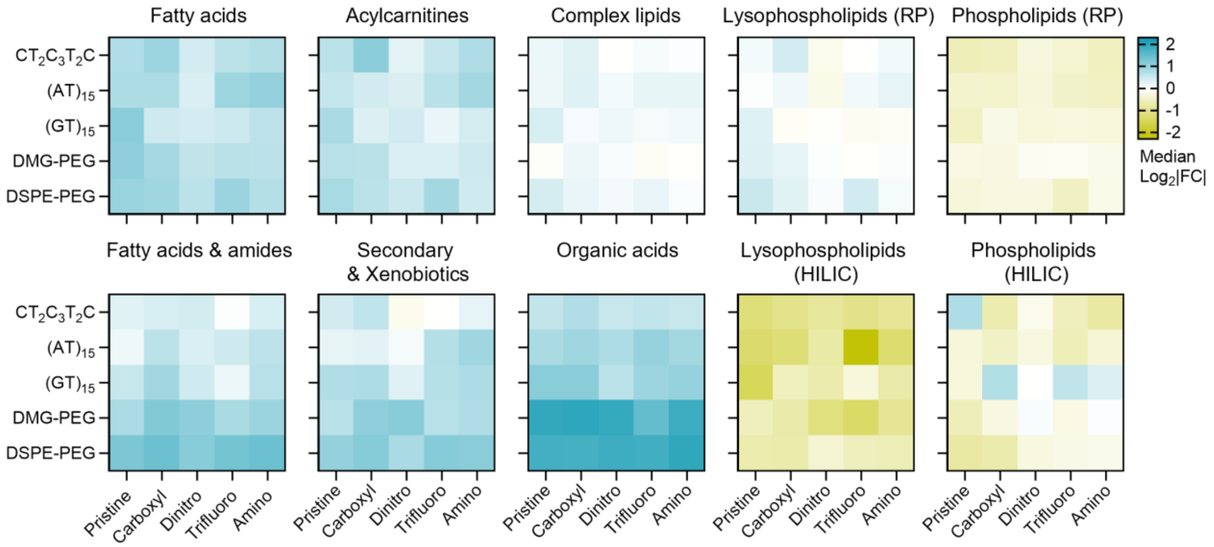

**Supplementary Figure 23. Polymer-QWD interaction heatmaps** for Track 1 metabolite classes across all nanotubes. Columns represent QWD types ordered by polarity (Pristine, Carboxyl, Dinitro, Trifluoro, Amino); rows represent polymer wrappings. Color bar indicates the enrichment (blue) and depletion (yellow) of metabolites based on the median $\log_2$FC (corona vs plasma). The top rows list metabolite classes detected in RP. The bottom rows list metabolite classes detected by HILIC.

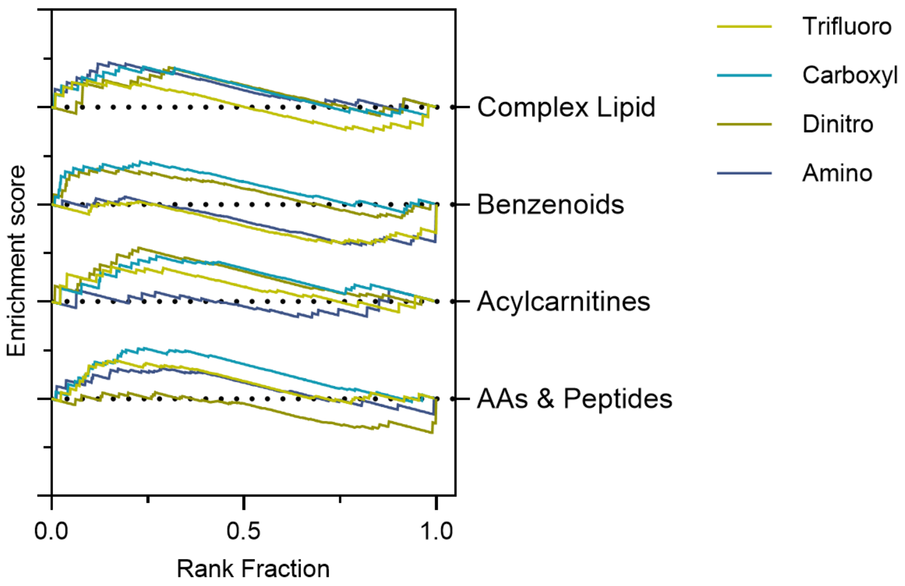

**Supplementary Figure** 24**.** Rank-ordered enrichment score of HILIC Track 2 classes across QWD types, pooled across all five polymer wrappings. Features are ranked by mean $\Delta log_2FC$ (QWD vs pristine). Most curves exhibit flat or irregular trajectories, indicating that Track 2 class members are broadly distributed within the ranked feature list rather than concentrated at enrichment extremes. Lysophospholipids and Acylcarnitines are the exceptions, showing modest QWD-dependent enrichment patterns described in the main text.

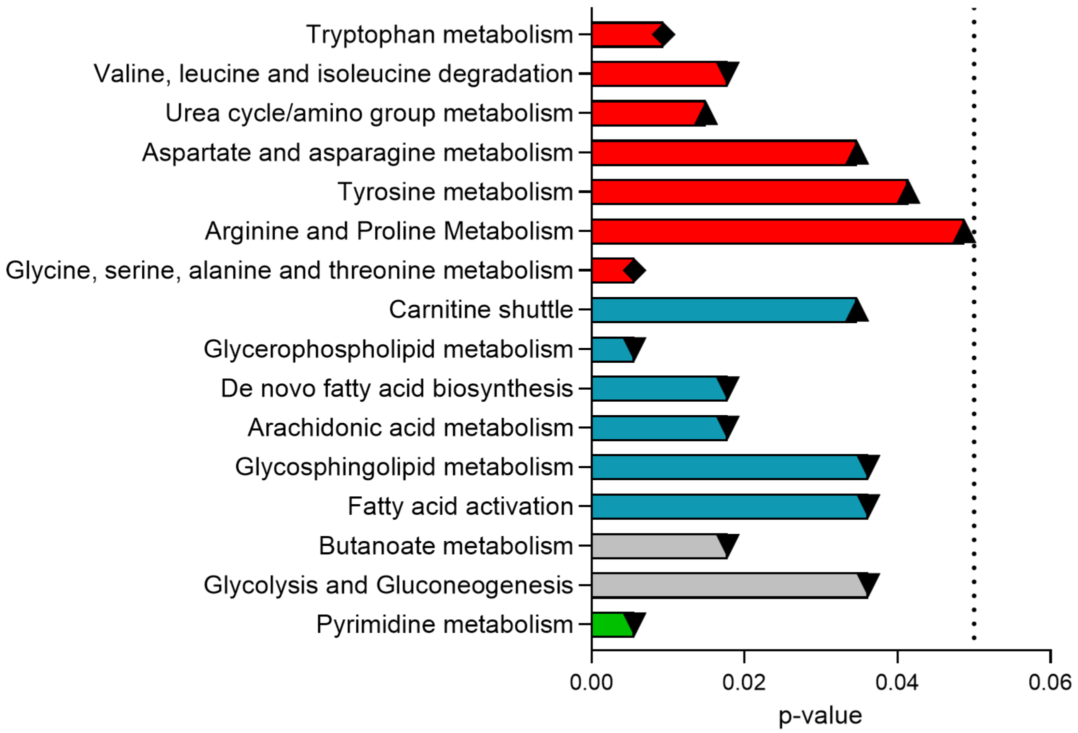

**Supplementary Figure 25. Mummichog pathway enrichment analysis of the HILIC metabolite corona.** Horizontal bars show -log(p-value) for pathways significantly enriched (permutation $p < 0.05$) among corona-enriched features pooled across all 25 nanotube conditions. Bar color indicates pathway category: Lipid metabolism (blue), amino acid metabolism (red), nucleotide metabolism (green), and other (gray). Triangles denote pathways detected in ESI+ (▲) or ESI- (▼) mode only; diamonds (◆) indicate pathways detected in both modes. Dashed line marks the significance threshold ($p = 0.05$).

**Supplementary Table 1.** Physicochemical descriptors of the four QWD aryl groups. Hammett σ values, computed from Hansch–Leo substituent constants, provide a measure of the electron-withdrawing (positive σ) or electron-donating (negative σ) character of each group. The McGowan volume and molar refractivity capture molecular size. Hydrodynamic radius ($R_h$) reflects the effective steric footprint of the defect in aqueous solution, and logP is the computed octanol–water partition coefficient.

| Abbreviation | QWD full name | logP | Charge at pH 7.4 | Hammett σ | $R_h$ (nm) | Mol. Refractivity (polarizability) |
|---|---|---|---|---|---|---|
| Carboxyl | 4-carboxylphenyl | −2.0 | −0.999 | 0.00 | 3.04 | 31.0 |
| Dinitro | 3,5-dinitrophenyl | +1.5 | 0.000 | 1.42 | 2.83 | 37.6 |
| Trifluoro | 3,4,5-trifluorophenyl | +2.6 | 0.000 | 0.74 | 2.44 | 26.1 |
| Amino | N,N-diethyl-4-aminophenyl | +3.7 | +0.126 | −0.83 | 3.48 | 49.8 |

**Supplementary Table 2.** IVW-combined $Log_2FC$ summary statistics. Per-condition IVW-combined $log_2FC$ statistics (number of features, median, mean, and percentage significant at $p < 0.05$ with $|log_2FC| > 1$) for all 25 nanotube conditions in both RP and HILIC modes, for the corona-versus-plasma and QWD-versus-pristine comparisons.

| Mode | IVW-combined features | Deduplicated IVW-combined features | Corona detected features | Median $log_2FC$ of all corona combined | Significant features with $\|log_2FC\|>1$ | % Significant |
|---|---|---|---|---|---|---|
| RP | 6453 | 6383 | 3943 | -0.0789 | 1098 | 27.8 |
| HILIC | 2732 | 2526 | 1037 | 0.236 | 478 | 46.1 |

| | | RP | | HILIC | |
|---|---|---|---|---|---|
| Polymer | QWD | n_feature | Median $log_2FC$ | n_feature | Median $log_2FC$ |
| $(AT)_{15}$ | Trifluoro | 2701 | -0.110 | 621 | 0.181 |
| | Carboxyl | 2490 | -0.071 | 619 | 0.108 |
| | Dinitro | 2253 | 0.040 | 574 | 0.307 |
| | Amine | 2322 | 0.010 | 552 | 0.458 |
| | Unfunctionalized | 2257 | -0.019 | 585 | 0.473 |
| $(GT)_{15}$ | Trifluoro | 2640 | -0.053 | 615 | 0.246 |
| | Carboxyl | 2165 | 0.103 | 553 | 0.527 |
| | Dinitro | 2707 | -0.044 | 577 | 0.700 |
| | Amine | 2915 | -0.091 | 507 | 0.490 |
| | Unfunctionalized | 2751 | -0.072 | 542 | 0.650 |
| $CT_2C_3T_2C$ | Trifluoro | 2683 | -0.142 | 654 | 0.017 |
| | Carboxyl | 2255 | -0.111 | 629 | 0.089 |
| | Dinitro | 2014 | 0.128 | 599 | 0.292 |
| | Amine | 2471 | -0.080 | 644 | -0.025 |
| | Unfunctionalized | 2271 | -0.055 | 653 | 0.082 |
| DMG-PEG | Trifluoro | 2361 | 0.027 | 486 | 1.098 |
| | Carboxyl | 1970 | 0.034 | 441 | 0.799 |
| | Dinitro | 2171 | 0.077 | 441 | 1.071 |
| | Amine | 2453 | -0.042 | 496 | 0.689 |
| | Unfunctionalized | 2309 | -0.040 | 443 | 0.662 |
| DSPE-PEG | Trifluoro | 2231 | 0.017 | 497 | 0.935 |
| | Carboxyl | 1888 | 0.248 | 472 | 1.026 |
| | Dinitro | 1952 | 0.143 | 448 | 1.172 |
| | Amine | 1796 | 0.144 | 531 | 1.095 |
| | Unfunctionalized | 2192 | 0.025 | 457 | 1.046 |

**Supplementary Table 3.** Metabolite class consolidation decisions. Mapping of original spectral library annotations to the consolidated class labels used for RP (5 classes) and HILIC (9 classes), with rationale based on chromatographic retention behavior, mass spectral properties, and biosynthetic relationships. md: Mass defect. m/z: mass to charge ratio.

**Reverse Phase**

| Consolidated Class | Constituent Classes | Rationale |
|---|---|---|
| Phospholipids | Glycerophospholipids (diacyl PC, PE, PI, PS, PG) | Two acyl chains; high m/z (>700); RP late elution |
| Complex Lipids | Glycerolipids, Sphingolipids, Sterol lipids | Neutral, highly hydrophobic; distinct m/z range |
| Lysophospholipids | Glycerophospholipids (lyso-species) | Single acyl chain; lower m/z than diacyl PL; separable by mass defect |
| Acylcarnitines | Acylcarnitines | Permanent positive charge; distinctive low mass defect |
| Fatty Acids | Fatty acids, Fatty acyls | Anionic at pH 7.4; early RP elution |

**HILIC**

| Consolidated Class | Constituent Classes | Rationale |
|---|---|---|
| Phospholipids | Glycerophospholipids (single-chain PE, PA) | Single-chain PL detected in HILIC; early HILIC RT Distinct m/z, md, and polarity fingerprint; the shared label preserves cross-mode consistency |
| Organic Acids | Organic acids, Bile acids, Carbohydrates | Small polar metabolites; anionic; md ~0.05; predominantly negative mode; early HILIC elution |
| Fatty Acids & Amides | Fatty acids, Fatty amides | Amphiphilic; intermediate HILIC RT |
| Secondary & Xenobiotic | Secondary metabolites, Xenobiotics, Sulfonates, Phenolics Terpenoids, Alkaloids, Polyphenols & Flavonoids, Vitamins & Cofactors | Diverse polar compounds; md ~0.10–0.15; m/z 150–300 Da; mixed polarity; overlapping aromatic/heterocyclic character late HILIC RT |
| Lysophospholipids | Lysophospholipids | Distinct m/z, md, and polarity fingerprint; the shared label preserves cross-mode consistency |
| Acylcarnitines | Acylcarnitines | Distinct m/z, md, and polarity fingerprint; the shared label preserves cross-mode consistency |
| Complex Lipids | Acylglycerols, Sphingolipids, Steroids & Bile Acids, other complex lipids | High md >0.25; nonpolar |
| Benzenoids | Benzenoids, Aromatic compounds | Low md ~0.076; m/z ~179 Da; 58% negative mode — distinct from Secondary & Xenobiotic |
| Amino Acids & Peptides | Amino acids, Peptides, Nucleobases, Indoles & Trp Metabolites, Sugars & Polyols, Nucleotides & Bases, Polar Metabolites | md ~0.05; m/z <200 Da; high polarity; early HILIC elution |

**Supplementary Table 4.** Per class test performance of the best-performing prediction models and Track1/2 assignment.

**Reverse Phase, Random Forest**

| Class | Precision | Recall | F1 | Track |
|---|---|---|---|---|
| Acylcarnitines | 1 | 1 | 1 | 1 |
| Complex Lipids | 0.9091 | 0.9259 | 0.9174 | 1 |
| Fatty Acids | 1 | 1 | 1 | 1 |
| Lysophospholipids | 1 | 1 | 1 | 1 |
| Phospholipids | 0.9333 | 0.918 | 0.9256 | 1 |
| Weighted | | | 0.9339 | |
| Cross validation | | | 0.9128 ± 0.014 | |

**HILIC, Random Forest**

| Class | Precision | Recall | F1 | Track |
|---|---|---|---|---|
| Acylcarnitines | 0.6667 | 0.3333 | 0.4444 | 2 |
| Amino Acids & Peptides | 0.3750 | 0.4839 | 0.4225 | 2 |
| Benzenoids | 0.4444 | 0.2353 | 0.3077 | 2 |
| Complex Lipids | 0.3333 | 0.2500 | 0.2857 | 2 |
| Fatty Acids & Amides | 0.5385 | 0.6176 | 0.5753 | 1 |
| Lysophospholipids | 0.6667 | 0.8571 | 0.7500 | 1 |
| Organic Acids | 0.5769 | 0.6383 | 0.6061 | 1 |
| Phospholipids | 0.5000 | 0.6667 | 0.5714 | 1 |
| Secondary & Xenobiotic | 0.5862 | 0.5000 | 0.5397 | 1 |
| Weighted | | | 0.5246 | |
| Cross validation | | | 0.4761 ± 0.020 | |

**Supplementary Table 5.** Feature deduplication summary per metabolite class. Number of features before and after name-based deduplication for each metabolite class in RP and HILIC, showing the number of collapsed adduct groups and percentage reduction. Acylcarnitines and phospholipids are the most affected classes due to the prevalence of sodium and potassium adducts.

**Reverse Phase**

| Class | n_full | n_dedup | n_collapsed | % Change |
|---|---|---|---|---|
| Phospholipids | 1708 | 1665 | 43 | -2.52 |
| Complex Lipids | 1917 | 1895 | 22 | -1.15 |
| Lysophospholipids | 2022 | 2018 | 4 | -0.20 |
| Acylcarnitines | 431 | 430 | 1 | -0.23 |
| Fatty Acids | 373 | 373 | 0 | 0 |
| **TOTAL** | 6453 | 6383 | 70 | -1.08 |

**HILIC**

| Class | n_full | n_dedup | n_collapsed | % Change |
|---|---|---|---|---|
| Phospholipids | 72 | 70 | 2 | -2.78 |
| Organic Acids | 642 | 592 | 50 | -7.79 |
| Fatty Acids & Amides | 247 | 224 | 23 | -9.31 |
| Secondary & Xenobiotic | 702 | 653 | 49 | -6.98 |
| Lysophospholipids | 130 | 127 | 3 | -2.31 |
| Acylcarnitines | 50 | 37 | 13 | -26.0 |
| Complex Lipids | 53 | 52 | 1 | -1.89 |
| Benzenoids | 105 | 86 | 19 | -18.1 |
| Amino Acids & Peptides | 192 | 149 | 43 | -22.4 |
| **TOTAL** | 2732 | 2529 | 203 | -7.43 |

**Supplementary Table 6.** Deduplication sensitivity analysis. Comparison of class-level enrichment statistics (median $\log_2$FC, interquartile range, direction, percentage of positive features) between the full feature set and the deduplicated feature set for all Track 1 classes across RP and HILIC. All class-level directions and statistical conclusions are preserved after deduplication.

**Reverse Phase**

| **Class** | Dataset | n_features | Median $\text{Log}_2$FC | q25 | q75 | % Positive | Direction |
|---|---|---|---|---|---|---|---|
| Phospholipids | Full | 1265 | -0.232 | -0.741 | 0.140 | 31.4 | Depleted |
| | Dedup | 1233 | -0.226 | -0.700 | 0.157 | 31.9 | Depleted |
| Complex Lipids | Full | 1262 | 0.063 | -0.381 | 0.672 | 52.9 | Enriched |
| | Dedup | 1245 | 0.058 | -0.373 | 0.667 | 52.7 | Enriched |
| Lysophospholipids | Full | 1065 | -0.110 | -0.804 | 0.471 | 43.6 | Depleted |
| | Dedup | 1062 | -0.108 | -0.789 | 0.471 | 43.7 | Depleted |
| Acylcarnitines | Full | 232 | 0.389 | -0.020 | 1.057 | 73.7 | Enriched |
| | Dedup | 231 | 0.393 | -0.012 | 1.058 | 74.0 | Enriched |
| Fatty Acids | Full | 170 | 0.428 | -0.037 | 1.178 | 73.5 | Enriched |
| | Dedup | 170 | 0.428 | -0.037 | 1.178 | 73.5 | Enriched |

**HILIC**

| Class | Dataset | n_features | Median $\text{Log}_2$FC | q25 | q75 | % Positive | Direction |
|---|---|---|---|---|---|---|---|
| Phospholipids | Full | 46 | -0.092 | -0.645 | 0.502 | 45.7 | Depleted |
| | Dedup | 45 | -0.102 | -0.638 | 0.503 | 44.4 | Depleted |
| Lysophospholipids | Full | 72 | -1.102 | -2.451 | -0.08 | 20.8 | Depleted |
| | Dedup | 70 | -1.088 | -2.446 | -0.075 | 21.4 | Depleted |
| Organic Acids | Full | 227 | 0.846 | -0.226 | 1.934 | 70 | Enriched |
| | Dedup | 213 | 0.846 | -0.233 | 1.913 | 69.5 | Enriched |
| Fatty Acids & Amides | Full | 134 | 0.601 | -0.238 | 1.543 | 69.4 | Enriched |
| | Dedup | 120 | 0.637 | -0.257 | 1.533 | 68.3 | Enriched |
| Secondary & Xenobiotic | Full | 285 | 0.14 | -1.024 | 1.337 | 53.3 | Enriched |
| | Dedup | 268 | 0.182 | -0.984 | 1.374 | 55.2 | Enriched |

**Supplementary Table 7.** Enrichment scores for HILIC Track 2 classes for pristine nanotubes. Enrichment scores (ES) for each Track 2 class across the five polymer wrappings (pristine nanotubes). Positive ES indicates class members concentrated at the QWD-enriched or corona-enriched end of the ranked feature list; negative ES indicates concentration at the depleted end.

| Polymer | Class | n_class | n_total | ES | Polymer enriched? |
|---|---|---|---|---|---|
| $(AT)_{15}$ | Acylcarnitines | 18 | 585 | -0.475 | N |
| | Complex Lipids | 14 | | 0.321 | Y |
| | Benzenoids | 27 | | -0.238 | N |
| | Amino Acids & Peptides | 28 | | -0.654 | N |
| $(GT)_{15}$ | Acylcarnitines | 19 | 542 | -0.521 | N |
| | Complex Lipids | 12 | | 0.446 | Y |
| | Benzenoids | 24 | | 0.331 | Y |
| | Amino Acids & Peptides | 27 | | -0.717 | N |
| $CT_2C_3T_2C$ | Acylcarnitines | 18 | 653 | -0.579 | N |
| | Complex Lipids | 16 | | -0.273 | N |
| | Benzenoids | 29 | | -0.273 | N |
| | Amino Acids & Peptides | 32 | | -0.649 | N |
| DMG-PEG | Acylcarnitines | 19 | 443 | -0.584 | N |
| | Complex Lipids | 8 | | 0.622 | Y |
| | Benzenoids | 20 | | 0.318 | Y |
| | Amino Acids & Peptides | 15 | | -0.617 | N |
| DSPE-PEG | Acylcarnitines | 16 | 457 | -0.356 | N |
| | Complex Lipids | 8 | | 0.654 | Y |
| | Benzenoids | 21 | | 0.460 | Y |
| | Amino Acids & Peptides | 20 | | -0.651 | N |

**Supplementary Table 8.** Enrichment scores for HILIC Track 2 classes across the four QWD types. Enrichment scores (ES) for each Track 2 class across the four QWD types (pooled across polymers). Positive ES indicates class members concentrated at the QWD-enriched or corona-enriched end of the ranked feature list; negative ES indicates concentration at the depleted end.

| QWD | Class | n_class | n_total | ES | QWD enriched? |
|---|---|---|---|---|---|
| Trifluoro | Acylcarnitines | 22 | 1032 | 0.359 | Y |
| | Complex Lipids | 23 | | 0.272 | Y |
| | Benzenoids | 40 | | -0.404 | N |
| | Amino Acids & Peptides | 36 | | 0.396 | Y |
| Carboxyl | Acylcarnitines | 22 | 1054 | 0.468 | Y |
| | Complex Lipids | 20 | | 0.417 | Y |
| | Benzenoids | 41 | | 0.439 | Y |
| | Amino Acids & Peptides | 37 | | 0.521 | Y |
| Dinitro | Acylcarnitines | 21 | 1026 | 0.554 | Y |
| | Complex Lipids | 21 | | 0.410 | Y |
| | Benzenoids | 40 | | 0.368 | Y |
| | Amino Acids & Peptides | 38 | | -0.351 | N |
| Amino | Acylcarnitines | 20 | 1073 | -0.163 | N |
| | Complex Lipids | 26 | | 0.458 | Y |
| | Benzenoids | 40 | | -0.419 | N |
| | Amino Acids & Peptides | 37 | | 0.316 | Y |

**Supplementary Table 9.** Mummichog pathway enrichment results. All pathways with permutation $p < 0.05$ from mummichog analysis of the HILIC dataset (positive and negative ionization modes combined), including pathway name, category, ESI mode, overlap size, pathway size, and p-value.

| Pathway | Category | ESI mode | Overlap | Pathway Size | p value |
|---|---|---|---|---|---|
| Glycerophospholipid metabolism | Lipid | NEG | 13 | 13 | 0.0056 |
| Glycine, serine, alanine & threonine metabolism | Amino acid | Both | 13 | 13 | 0.0056 |
| Pyrimidine metabolism | Nucleotide | NEG | 13 | 13 | 0.0056 |
| Tryptophan metabolism | Amino acid | Both | 12 | 12 | 0.0094 |
| Urea cycle / amino group metabolism | Amino acid | POS | 13 | 13 | 0.015 |
| De novo fatty acid biosynthesis | Lipid | NEG | 10 | 10 | 0.0178 |
| Valine, leucine & isoleucine degradation | Amino acid | NEG | 10 | 10 | 0.0178 |
| Butanoate metabolism | Other | NEG | 10 | 10 | 0.0178 |
| Arachidonic acid metabolism | Lipid | NEG | 10 | 10 | 0.0178 |
| Aspartate & asparagine metabolism | Amino acid | POS | 10 | 10 | 0.0347 |
| Carnitine shuttle | Lipid | POS | 10 | 10 | 0.0347 |
| Glycolysis & Gluconeogenesis | Other | NEG | 8 | 8 | 0.0362 |
| Glycosphingolipid metabolism | Lipid | NEG | 8 | 8 | 0.0362 |
| Fatty acid activation | Lipid | NEG | 8 | 8 | 0.0362 |
| Tyrosine metabolism | Amino acid | POS | 9 | 9 | 0.0414 |
| Arginine & Proline metabolism | Amino acid | POS | 8 | 8 | 0.0487 |
